# Characterization of the interactive effects of labile and recalcitrant organic matter on microbial growth and metabolism

**DOI:** 10.1101/524777

**Authors:** Lauren N. M. Quigley, Abigail Edwards, Andrew D. Steen, Alison Buchan

## Abstract

Geochemical models typically represent organic matter (OM) as consisting of multiple, independent pools of compounds, each accessed by microorganisms at different rates. However, recent findings indicate that organic compounds can interact within microbial metabolisms. The relevance of interactive effects within marine systems is debated and a mechanistic understanding of its complexities, including microbe-substrate relationships, is lacking. As a first step toward uncovering mediating processes, the interactive effects of distinct pools of OM on the growth and respiration of marine bacteria, individual strains and a simple, constructed community of Roseobacter lineage members were tested. Isolates were provided with natural organic matter (NOM) and different concentrations (1, 4, 40, 400 μM-C) and forms of labile organic matter (acetate, casamino acids, tryptone, coumarate). The microbial response to the mixed substrate regimes was assessed using viable counts and respiration in two separate experiments. Two marine bacteria and a six-member constructed community were assayed with these experiments. Both synergistic and antagonistic growth responses were evident for all strains, but all were transient. The specific substrate conditions promoting a response, and the direction of that response, varied amongst species. These findings indicate that the substrate conditions that result in OM interactive effects are both transient and species-specific and thus influenced by both the composition and metabolic potential of a microbial community.

## Introduction

2.5 Tg-C of terrestrially-derived dissolved organic matter (t-DOM) flows through riverine systems annually, where the microbial community preferentially utilizes the more labile components (Vannote *et al.* 1980; Hedges, Keil and Benner 1997). This process leads to the development of an increasingly recalcitrant organic carbon pool, enriched in aromatic moieties, as headwaters move towards coastal margins (Sun *et al.* 1997; Mannino and Harvey 2000). Most chemical tracers diagnostic of t-DOM (e.g. lignin-derived phenols) are removed before reaching the open oceans (Hedges, Keil and Benner 1997; Osburn *et al.* 2016), suggesting that this material is transformed in land-sea margins. Microbial degradation clearly contributes to the disappearance of t-DOM in these dynamic aquatic systems (Ward *et al.* 2013).

It has recently been postulated that biological interactions with different pools of organic compounds drive OM transformations in aquatic environments (Guenet *et al.* 2010; Bianchi 2011). This hypothesis has been framed within the concept of the priming effect (PE). Under the broadest definition of the term, PE occurs when the addition of a labile carbon substrate and/or nutrients alters the rate at which microorganisms degrade recalcitrant organic carbon (Kuzyakov, Friedel and Stahr 2000). These interactive effects are non-additive and can be either positive (synergistic) or negative (antagonistic). The microbial response may rely critically on the concentration and molecular composition of organic compounds, experimental timescale, nutrient status and microbial community composition (Blagodatskaya and Kuzyakov 2008; Catalán *et al.* 2015a; Steen, Quigley and Buchan 2016). PE has long been recognized as an important factor in soil organic matter turnover. However, this framework has only recently been applied to aquatic systems where its present role is enigmatic (Jenkinson, Fox and Rayner 1985; Guenet *et al.* 2010; Bianchi 2011). Bengtsson et al. posit the variable PE responses reported in the aquatic sciences literature suggests OM interactive effects are likely context dependent. As such, an improved mechanistic understanding of the microbial response to mixed OM pools is needed to enable predictive modeling of OM fate in various environments (Bengtsson, Attermeyer and Catalán 2018).

The salt marshes fringing the coast of the Southeastern United States, and the microbial communities residing within these systems, provide a relevant system in which to study factors relevant to OM interactions and microbial processing. The rivers flowing through these marshes carry 400 to 2300 μM-C dissolved organic carbon (DOC), approximately 75% of which is terrestrially-derived (Alberts and Takács 1999). Additionally, these salt marshes are among the most productive ecosystems on Earth, with net primary production rates ranging from 0.2 to 2.25 kg C m^−2^ yr^−1^ (Wiegart and Freeman 1990; Hyndes *et al.* 2014). Within these systems, autochtonous labile inputs, from salt marsh vegetation and phytoplankton, mix with the recalcitrant t-DOM imported by riverine systems at the land-sea interface, setting the stage for OM interactions that may stimulate resident coastal microbial communities to degrade recalcitrant t-DOM. Potential for positive, albeit transient, priming of Southeastern US coastal microbial communities has been demonstrated (Steen, Quigley and Buchan 2016). However, the specific factors that control OM interactive effects at the level of individual environmentally relevant bacteria and/or communities of bacteria, have not been elucidated.

Members of the Roseobacter clade of marine bacteria are among the most numerically abundant and active members of the coastal bacterial communities, and several representative strains have been isolated from Southeastern US estuaries (e.g. Gonzalez *et al.* 1997; González, Kiene and Moran 1999; Slightom and Buchan 2009). Success of the lineage has largely been attributed to metabolic diversity, including growth on a wide range of plant-derived aromatic compounds characteristic of t-DOM (Moran *et al.* 2007; Mou *et al.* 2008; Medeiros *et al.* 2017; Sipler *et al.* 2017). Growth assays are supported by genome analyses which indicate Roseobacters often possess multiple catabolic pathways for aromatic compound degradation (Newton *et al.* 2010). Given their abundance, metabolic activity, and ability to oxidize plant-derived aromatic monomers, members of the Roseobacter clade are ideal lab cultivars to examine how representative members of the estuarine community may undergo interactive effects to degrade t-DOM.

Here we assess the influence of labile carbon concentration and chemical identity on the growth dynamics and respiration of representative marine bacteria provided an environmentally relevant and natural source of organic matter (NOM). The NOM utilized in these experiments was derived from the Suwanee River, a Southeastern US blackwater river, and is enriched in aromatic moieties of lignin origin (Her *et al.* 2003). We used monocultures of two coastal Roseobacter species, *Sagittula stellata* E-37 and *Citreicella* sp. SE45, both of which were isolated from Southeastern coastal waters and have demonstrated abilities to degrade plant-derived recalcitrant compounds (Gonzalez *et al.* 1997; Frank 2016; Chua 2018). We constructed community of six coastal bacteria that included these two strains community as well as four other Roseobacter strains selected based on the number (1-6) and types of aromatic carbon catabolism pathways present in their genomes (Table 1). In this study, 16 labile organic matter (LOM) conditions were tested in a fully factorial experiment. Four substrates, ranging from simple to chemically complex (sodium acetate, coumarate, casamino acids + tryptophan, and tryptone, in order of increasing chemical complexity) at four concentrations (1, 4, 40, and 400 μM-C; Table 2) were provided as growth substrates for monocultures of *S. stellata* and *Citreicella* sp. SE45 and the six-member constructed community. Sources of LOM were chosen to represent a gradient of chemical complexities that are differentially processed by microbes: sodium acetate and casamino acids + tryptophan are likely shunted directly into central metabolism; tryptone is a mixture of oligo-peptides, many of which require initial extracellular enzymatic breakdown before products are transported across the cell membrane and enter central metabolism; and coumarate, an aromatic monomer derived from lignin (Hedges *et al.* 1988). Cleavage of the aromatic ring requires specific pathways that are found in a limited number of microbes and are most often subject to catabolite repression (Dal, Steiner and Gerischer 2002; Mazzoli *et al.* 2007). Of the six bacterial isolates tested, only *S. stellata* and *Citreicella* sp. SE45 possess the ability use coumarate as a sole carbon source.

**Table 1.**
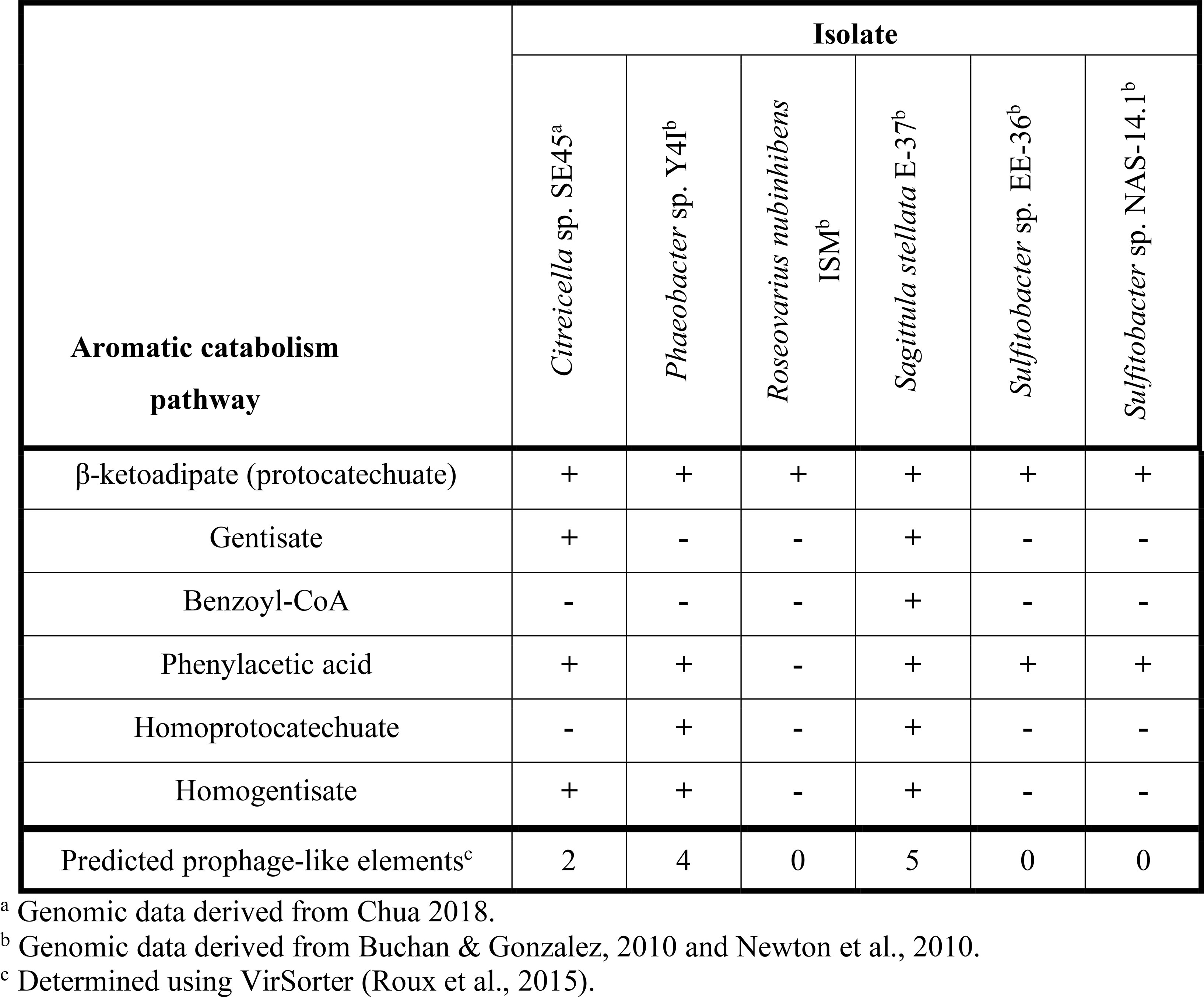
Genomic evidence for aromatic carbon catabolism pathways and prophages present in Roseobacter strains used in this study.

**Table 2.** Organic carbon composition of the comparative treatments groups used to test for interactive effects.

| Comparative treatment group | NoC<sup>a</sup> | LOM<sup>b</sup> | NOM<sup>c</sup> | mix<sup>d</sup> |
| --- | --- | --- | --- | --- |
| 400 μM-C | No OC added | 400 μM-C LOM | 2 mM-C NOM | 400 μM-C LOM (16.67%)<br>2 mM-C NOM (83.33%) |
| 40 μM-C | No OC added | 40 μM-C LOM | 2 mM-C NOM | 40 μM-C LOM (1.67%)<br>2 mM-C NOM (98.33%) |
| 4 μM-C | No OC added | 4 μM-C LOM | 2 mM-C NOM | 4 μM-C LOM (0.17%)<br>2 mM-C NOM (99.83%) |
| 1 μM-C | No OC added | 1 μM-C LOM | 2 mM-C NOM | 1 μM-C LOM (0.04%)<br>2 mM-C NOM (99.96%) |

<sup>a</sup> No carbon added (base medium), tests for microbial activity in the absence of added organic carbon.
<sup>b</sup> Labile organic matter, (acetate, casamino acids + tryptophan, coumarate, or tryptone) added at one of four concentrations.
<sup>c</sup> Natural organic matter (recalcitrant organic matter).
<sup>d</sup> Mixed substrate treatments. Values in parentheses indicate the relative contribution of LOM and NOM to the total organic carbon pool. To assess interactivity in mix treatments, those data are compared to composite data which is the sum of results of the LOM and NOM treatments.

## Materials and Methods

### Strains, media and growth conditions

*Sagittula stellata* sp. E-37, *Citreicella* sp. SE45, *Phaeobacter* sp. Y4I, *Roseovarius nubinhibens* ISM, *Sulfitobacter* sp. EE-36, and *Sulfitobacter* sp. NAS-14.1 were routinely grown on an aromatic basal medium (ABM) containing per liter 8.7 μM KCl, 8.7 μM CaCl_2_, 43.5 μM MgSO_4_, and 174 μM NaCl with 225 nM K_2_HPO_4_, 13.35 μM NH_4_Cl, 71 mM Tris-HCl (pH 7.5), 68 μM Fe-EDTA, trace metals (7.8492 mM Nitroloacetic acid, 0.5325 mM MnSO_4_ *H_2_ O, 0.4203 mM CoCl_2_ *6H_2_ O, 0.3478 mM ZnSO_4_ *7H_2_ O, 0.0376 mM CuSO_4_, 0.1052 mM NiCl_2_ *6H_2_ 0, 1.1565 mM Na_2_ SeO_3_, 0.4134 mM Na_2_ MoO_4_ *2H_2_ O, 0.3259 mM Na_2_WO_4_*2H_2_O, 0.2463 mM Na_2_SiO_3_*9H_2_O) and trace vitamins (0.0020% vitamin H [Biotin)], 0.0020% folic acid, 0.0100% pyridoxine-HCl (B6), 0.0050% riboflavin (B2), 0.0050% thiamine (B1), 0.0050% nicotinc acid, 0.0050% pantothenic acid (B5), 0.0001% cyanocobalamin (B12), 0.0050% *p*-aminobenzoic acid). These strains were routinely passaged on ABM containing 10 mM sodium acetate. Four of these strains (E-37, SE45, Y4I, and EE-36) were isolated from Southeastern US coastal waters, while NAS-14.1 was isolated from North Atlantic off-shore waters and ISM from the Caribbean Sea (Buchan *et al.* 2000; Cude *et al.* 2012). The bacteria were routinely cultured at 30°C, shaking, in the dark. This temperature condition is nominally representative of Southeastern US salt marshes which are tidally influenced and where average water temperatures are close to 30°C from June through September (The Southeast Reagional Climate Center, University of North Carolina, Chapel Hill, NC). Suwannee River natural organic matter (NOM), obtained from the International Humic Substance Society (IHSS, St. Paul, MN) was used as a representative t-DOM. This material is a discipline standard for natural organic matter (Her *et al.* 2003). Incubations occurred in the dark as the aromatic moieties in NOM are sensitive to photodegradation. NOM is provided in lyophilized form from IHSS and was suspended in Milli-Q water and 0.22 *μ*m filter-sterilized prior to addition to the medium. NOM was held at a constant concentration of 2 mM-C for all experiments. ^13^C NMR estimates of carbon distribution provided by IHSS show that Suwannee NOM comprised of roughly 25% aromatic residues.

Four different forms of labile organic matter (LOM) (sodium acetate, casamino acids + tryptophan, coumarate, and tryptone) were added at four concentrations (400, 40, 4, and 1 μM-C) using ABM as the base medium. Cultures were grown for 14 days in the dark at 30°C, with shaking. Substrate concentrations were selected after a preliminary experimentation using a LOM concentration gradient of 400 μM-C to 20 nM-C. All glassware used was combusted at 450° C for at least four hours to remove trace organic carbon. All experiments utilized cultures preconditioned on 2 mM-C *p*-hydroxybenzoic acid to match the carbon concentration of the Suwannee River NOM utilized in the mesocosms. The growth rates of all strains on this substrate at this concentration are comparable and mid-exponential phase cultures were used as inoculum at volumes of 10 – 100 μl. As cells were not washed prior to transfer to fresh media, there may have been some modest carryover of *p*-hydroxybenzoate (< 2μM). Nonetheless, carryover would have been consistent across treatments for a given strain or the six-member community and comparisons were always made to composite data (NOM plus LOM alone controls) which would have the same amount of carryover. Viable cell density experiments were carried out in volumes of 10 mL while respirometer incubations were in 125 mL volumes.

### Experimental treatments

All experiments assessed interactive effects of organic matter by comparing viable cell density or respiration in a treatment containing both labile and recalcitrant organic matter to the sum of growth or respiration in treatments containing only one of those carbon sources. There were a total of four treatments: NoC (no carbon addition control), LOM (labile organic matter alone), NOM (Suwannee River natural organic matter alone), and “mix” (LOM + NOM treatments) (Table 2). The NoC controls lacked both LOM and NOM, serving as a control for bacterial growth on medium components. The LOM treatment consisted of LOM under the same conditions as the corresponding mix treatment. The NOM treatment contained 2 mM-C Suwannee River NOM as the sole carbon source. The mix treatment had both 2 mM-C NOM and one of four concentrations of the different LOMs. Composite was calculated by adding the response of LOM alone and NOM alone. The microbial seeding density for all experiments was ~ 1×10^4^ cells mL^−1^, cell densities are reported in figures and tables. For the constructed community inoculum, equal representation of each strain was targeted.

For each treatment, viable cell abundance and community composition were measured. As we were motivated to understand the ability of different OM mixtures to support the growth of marine bacteria, viable counts were monitored rather than direct microscopic counts or DNA-based approaches, which generally do not readily distinguish between living and dead cells. Viable counts have the additional advantage over direct counts that it is easy to distinguish between individual Roseobacter strains (see below). Viable counts were obtained by serial dilution in ABM. Dilutions were plated on YTSS agar, a complex medium (per liter: 5 g yeast extract, 2.5 g tryptone and 15 g sea salts) and incubated in the dark at 30°C. Single strain plates were incubated for two days, while constructed community plates were incubated for four days in order to allow the development of identifying pigment. Plates with 30-300 colonies were counted. Cultures were spot-checked for cell clumping by microscopy and none was evident (Fig. S1).

Due to the impraticality of obtaining all of the necessary samples from a single set of experimental samples, two parallel sets of the same experiment were performed. A set of incubations for viable counts was first performed and the results from those incubations were used to inform the conditions selected for incubation in a respirometer. For the cell abundance and community composition experiment, culture aliquots were collected on days 0, 1, 2, 4, 7, 10 and 14. Community composition was determined by colony morphology, as each strain of Roseobacter in the community had a unique, readily identifiable, colony morphology (Fig. S2). Respiration was monitored in a separate set of microcosms using a Micro-Oxymax respirometer (Columbus Instruments, Columbus, OH), in which cumulative CO_2_ production was measured by infrared absorbance continuously throughout a shorter (7 day) incubation period.

### Data Analysis

To assess interactive effects of mixed substrate treatments, the sum of the viable cell density or CO_2_ production in the LOM and NOM treatments was calculated and termed “composite”, which represents the case in which growth on LOM and NOM are independent. The timing, extent and nature (synergistic or antagonistic) of interactive responses was determined through comparision of the resulting composite data and that from the mixed substrate treatments.

All data analysis was performed using the R statistical platform and visualized using the ggplot2 package (Wickham 2009; R Core team 2015). Raw data and scripts are posted at http://github.com/lnmquigley/roseo_priming_2018. Cell densities were log-transformed and sub-setted by day. For each day, a three-way ANOVA was performed to determine whether differences in cell densities were being driven by treatment, concentration or source of LOM. Because the experimental design was unbalanced, two three-way ANOVAs were performed on the final time point in the respirometer incubations in order to determine the factors influencing CO_2_ accumulation. Additionally, rates were calculated during exponential CO_2_ production, and three-way ANOVAs were employed to identify factors influencing the rate of CO_2_ production. For all ANOVAs, Fisher’s least significant difference was used as a post hoc test and *p*-values were adjusted to correct for the false discovery rate using the Benjamini-Hochberg correction (Benjamini and Hochberg 1995).

Community composition was determined by visually identifying constructed community members based on their distinct colony morphologies (Fig. S2). In order to calculate α diversity in the constructed community experiments, Shannon entropy was calculated for each culture, which was then exponentially transformed into Hill numbers, also known as effective species number (Jost 2007). A three-way ANOVA was performed to determine the relationship between effective species number and treatment, concentration and source of LOM. The *p*-values obtained from the Fisher’s least significant difference were adjusted using the Benjamini-Hochberg correction to account for multiple comparisons. A Bray-Curtis dissimilarity matrix was calculated using all constructed community cultures for each day. In order to determine sources of variation (treatment, concentration, and/or source of LOM) within the Bray-Curtis dissimilarity matrix, a permutational MANOVA was employed using the Adonis function in the R package vegan (Oksanen *et al.* 2017).

## Results

### Substrate preferences vary between individual strains

To assess the extent to which each LOM type and concentration could support the growth of the tested coastal marine bacteria, we monitored viable counts of monocultures of E-37 and SE45 as a function of organic matter treatment. Viable cell abundances for SE45 and E-37 increased two to three orders of magnitude within the first 24 hours of incubation, depending on the concentration of LOM provided (Fig. 1). In both strains, LOM type and concentration interacted significantly to drive cell densities at each time point (three-way ANOVA, n=5, *p*<0.001, Tables S1 and S2), with the single exception of E-37 on Day 14 (Table S2). For all four LOM types, the two lowest concentrations of LOM (1 and 4 μM) did not support reliable growth, relative to No C added controls, of either of the two monocultures over the course of the experiment (Fig. 1). With the exception of E-37 provided 40 μM tryptone, neither of the two bacterial isolates showed consistently robust growth at 40 μM with the remaining LOM substrates (Fig. 1). For all LOM types, the highest concentration of labile carbon (400 μM) showed significantly enhanced growth of both the strains (7-15 times greater than No C). A general trend emerged for all cultures in which cell viability increased rapidly at the start of the experiment and was followed by a decline beginning at Day 4 or later. SE45 saw declines ranging from 73-85% of viable cells in all of the 400 μM LOM alone treatments. However, viable cells remained significantly (at least 3-fold) higher than No C controls throughout the course of the experiment (Fig. 1). In contrast, E-37 demonstrated a more rapid decline in viability. By the end of the 14-day incubation period, strain E-37 had < 1% of maximum viable cells remaining in the LOM alone treatments provided 400 μM-C acetate, casamino acids and coumarate. Furthermore, viable counts in those cultures were indistinguishable from No C controls by Day 10 or earlier. Viable count data indicate that both monocultures and the community are able to use a small fraction of NOM-derived carbon in the absence of any LOM (Fig. 1).

**Figure 1.**
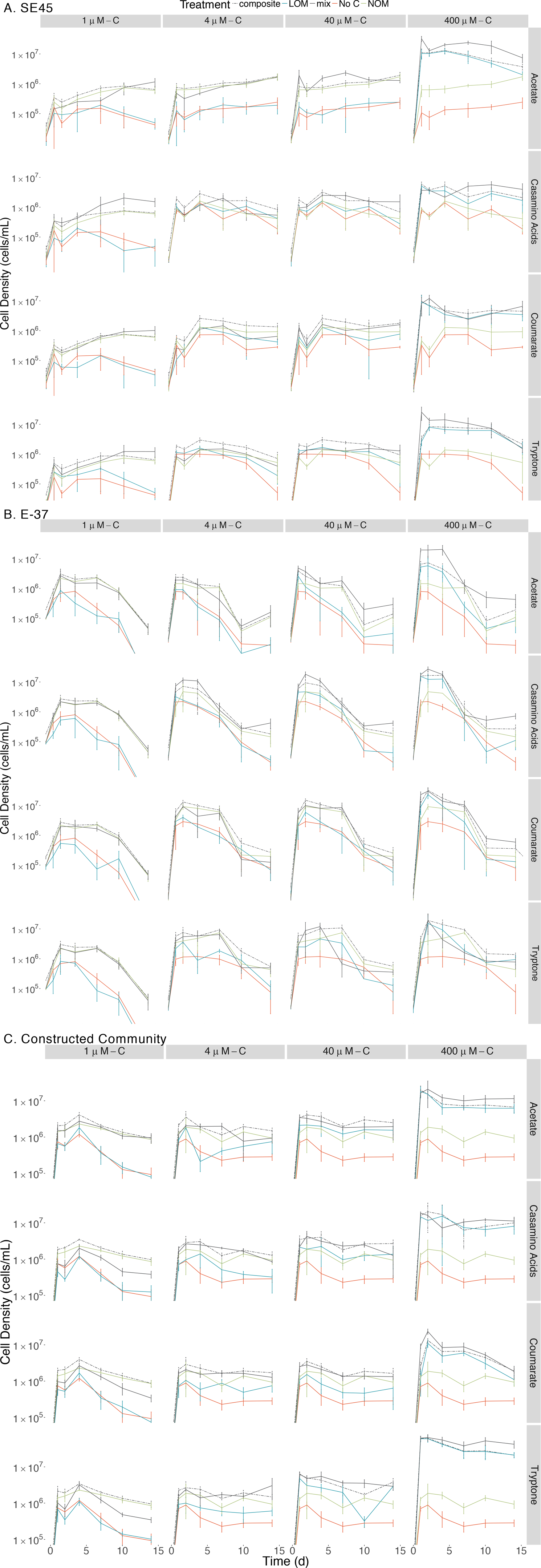
Viable counts for monocultures of (A) SE45, (B) E-37 and (C) constructed communities in composite (dashed line), LOM alone (blue line), mix (black line), No Carbon control (red line) and NOM alone control (green line) treatments. The composite treatment is the sum of results of the LOM and NOM treatments. Points represent the mean (n=3-5); error bars represent one standard deviation from the mean. Seeding densities for SE45, E-37 and six-member constructed community were 1.51 × 10^4^ CFU/mL (±5.1 × 10^3^), 4.23 × 10^4^ CFU/mL (±9 × 10^3^) and 7.01 × 10^3^ CFU/mL (±2.6 × 10^3^), respectively. Significant interactive effect for individual timepoints are shown in Fig. S3 (E-37 and SE45) and Fig. S4 (constructed community).

Each strain demonstrated unique and apparent preferences for the four different LOM types. Given the boom and bust growth dynamics described above, we focused on maximal viable counts within the first 48 hrs of the experiment for all LOM types provided at the highest concentration (400 μM). E-37 reached the highest cell densities on coumarate, nearly ten-fold higher viable counts compared to No C controls (2.3×10^7^ ± 6.28×10^6^ vs 2.86×10^6^ ± 9.34×10^5^ CFU/mL) and lowest on acetate (5.18×10^6^ ± 1.58×10^6^ CFU/mL). E-37 grew equally well on casamino acids and tryptone. SE45 growth was equivalent on all substrates except casamino acids, for which its viable counts were ~50% of those grown on the other three substrates within the first few days of the experiment (Fig. 1).

### Individual strains show differential responses to mixed organic matter treatments

For the mixed substrate experiments, NOM was held at a constant concentration of 2 mM-C, consistent with OC concentrations in Georgia coastal estuaries (Alberts and Takács 1999). To assess interactive growth responses, mixed substrate treatments (mix), which included a source of LOM and NOM in the same treatment, were compared to a composite class of data: the additive response of the LOM alone and NOM alone treatments. This allowed us to assess synergistic or antagonistic interactions of LOM and NOM on bacterial growth in the mixed treatments.

The individual strains displayed differing response to the various treatments. SE45 reached the highest viable cell densities in the mix treatments (LOM + NOM) with the highest LOM concentrations (400 μM-C; Fig. 1). Final viable cell densities increased with increasing LOM concentrations. While E-37 viable cell densities generally tracked with LOM concentrations, the differences in maximum cell densities across LOM type and concentration were less than an order of magnitude, compared to on average 10-fold difference in SE45 treatments between 400 μM-C and the lower concentrations.

While both strains displayed a significant growth response to the majority (≥75%) of mixed-substrate treatments, the effect was always transient (Fig. 1, Table 3). Both synergistic (positive) and antagnostic (negative) responses were observed, and the responses were species-specific. A significant synergistic response was seen for SE45 on all four LOM substrates at the highest concentration (400 μM-C). However, this was displayed at different time points for the different LOMs (Fig. S3, Table 3). Conversely, antagonistic interactions (i.e., composite cell densities significantly higher than those in mixtures) were observed for all LOM types at 4 μM-C (Fig. S3, Table 3) with this strain. Inconsistent trends were observed in other LOM concentration treatments. E-37 also displayed a significant response to all four LOM substrates at the highest concentrations, but the effect was negative on one substrate: tryptone (Fig. S3, Table 3). Growth of E-37 was negatively influenced at some point during the experiment for all concentrations of tryptone, except the lowest (1 μM–C). While a synergistic response was observed with coumarate at the highest concentration, antagonistic responses were observed with this substrate at the three lower concentrations. When E-37 displayed a significant growth response on casamino acids, it was always synergistic.

**Table 3.**
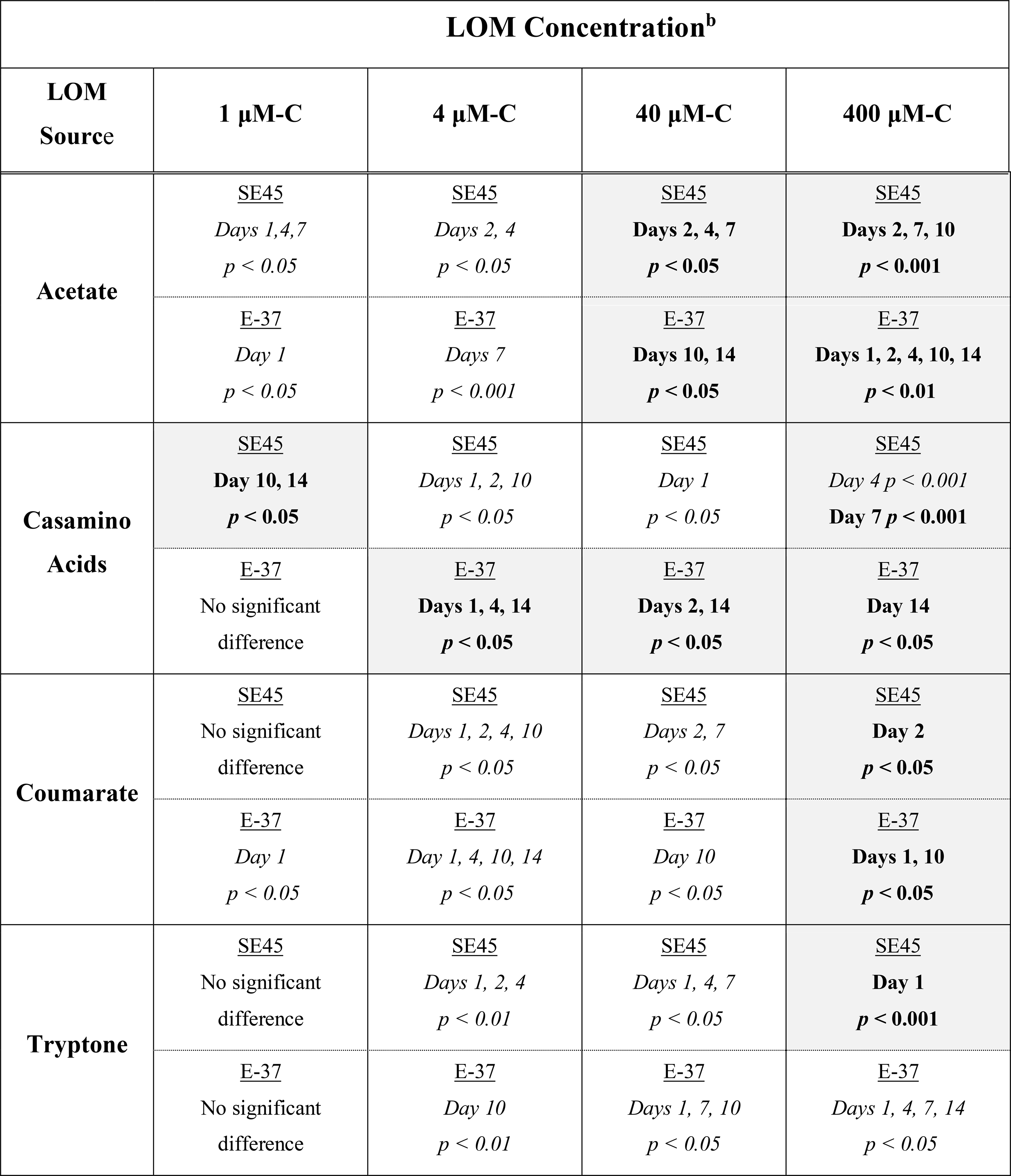
Probability values^a^ for monoculture interactive effects experiments shown in Figure 1 for strains SE45 and E-37.

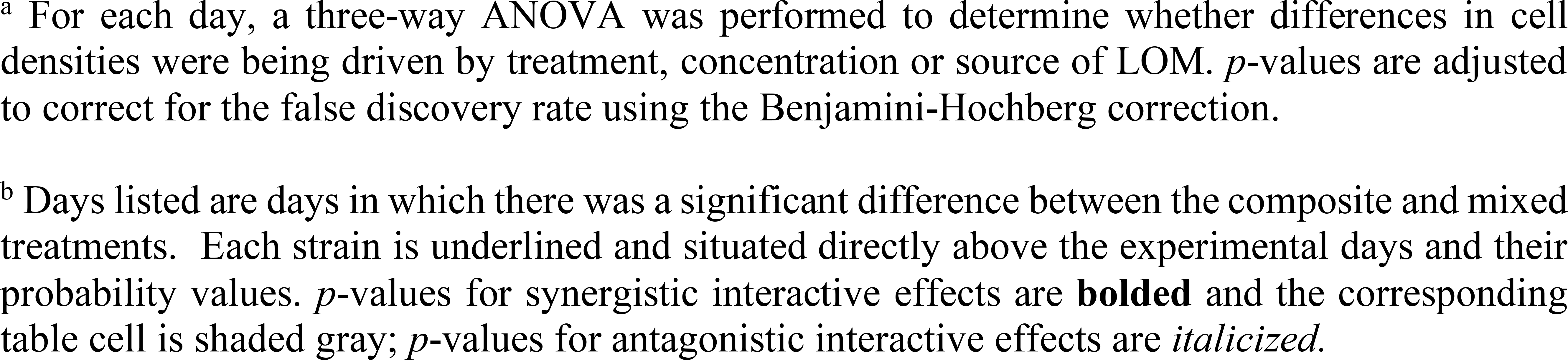

Due to the differential growth responses of the two strains to different concentrations of casamino acids, additional experiments were performed to monitor respiration at all concentrations of this LOM. Respiration assays were also performed on cultures provided acetate and coumarate at the highest LOM concentration (400 μM-C) to provide comparative information on the influence of different chemical compositions of LOM to microbial metabolism. These assays were run for seven days as the majority of culture growth from the previous experiment occurred within the first few days of incubation (Fig. 1). As a result of automated sampling, the respiration data provided higher temporal resolution. However, due to the sensitive nature of the probes it was neither possible to agitate the culture vessels nor take samples for viable counts throughout the course of the incubation, as was done for the previously described experiment. Instead, viable counts were performed for the seeding inoculums and each vessel at the final (Day 7) time points (Table S3). These values likely do not reflect the maximum viability of these cultures which is anticipated to have occurred earlier in the experiment, consistent with what was observed for the first experiment (see Fig. 1). Indeed, it is likely that by Day 7, cultures would be experiencing a decline in cell viability for several of the treatments.

The response by SE45 to mixed conditions when measured via respiration matched the results found in the initial experiment for 1, 4, and 400 μM-C casamino acids (Fig. 2). However, mixed and composite CO_2_ production were statistically indistinguishable from each other with cultures provided 400 μM-C coumarate and 40 μM-C casamino acids (Fig. 2, Table 5), despite the fact that these treatments exhibited significant synergistic and antagonistic responses, respectively, when assayed by viable count during the initial experiment (Fig. 1). CO_2_ production in NOM alone treatments was statistically indistinguishable from mixed OM treatments when SE45 was provided low concentrations of casamino acids as well as the highest (1, 4, and 400 μM; Student’s t-test *p* > 0.05). However, CO_2_ production on NOM was significantly lower than the mixed treatments at 40 μM-C casamino acids (Student’s t-test *p* < 0.05) and acetate and coumarate at 400 μM-C (Student’s t-test *p* < 0.05). At low concentrations of casamino acids, the mixed and composite treatments of E-37 were statistically indistinguishable according to the ANOVA model used (Table 5). The low concentration mixed treatments for E-37 all had significantly higher rates of CO_2_ production (~2-10 fold) than their corresponding LOM alone treatments (Tables S5-7). E-37 produced a synergestic response when stimulated with 400 μM-C acetate, casamino acids and coumarate, yielding cumulative CO_2_ production values that were 2-to 5-fold higher than the corresponding composite data (the sum of the NoC and LOM alone treatments) and rates that were 3-to 10-fold higher than the corresponding LOM alone data. (Fig. 2B, Tables 5, S5-7).

**Figure 2.**
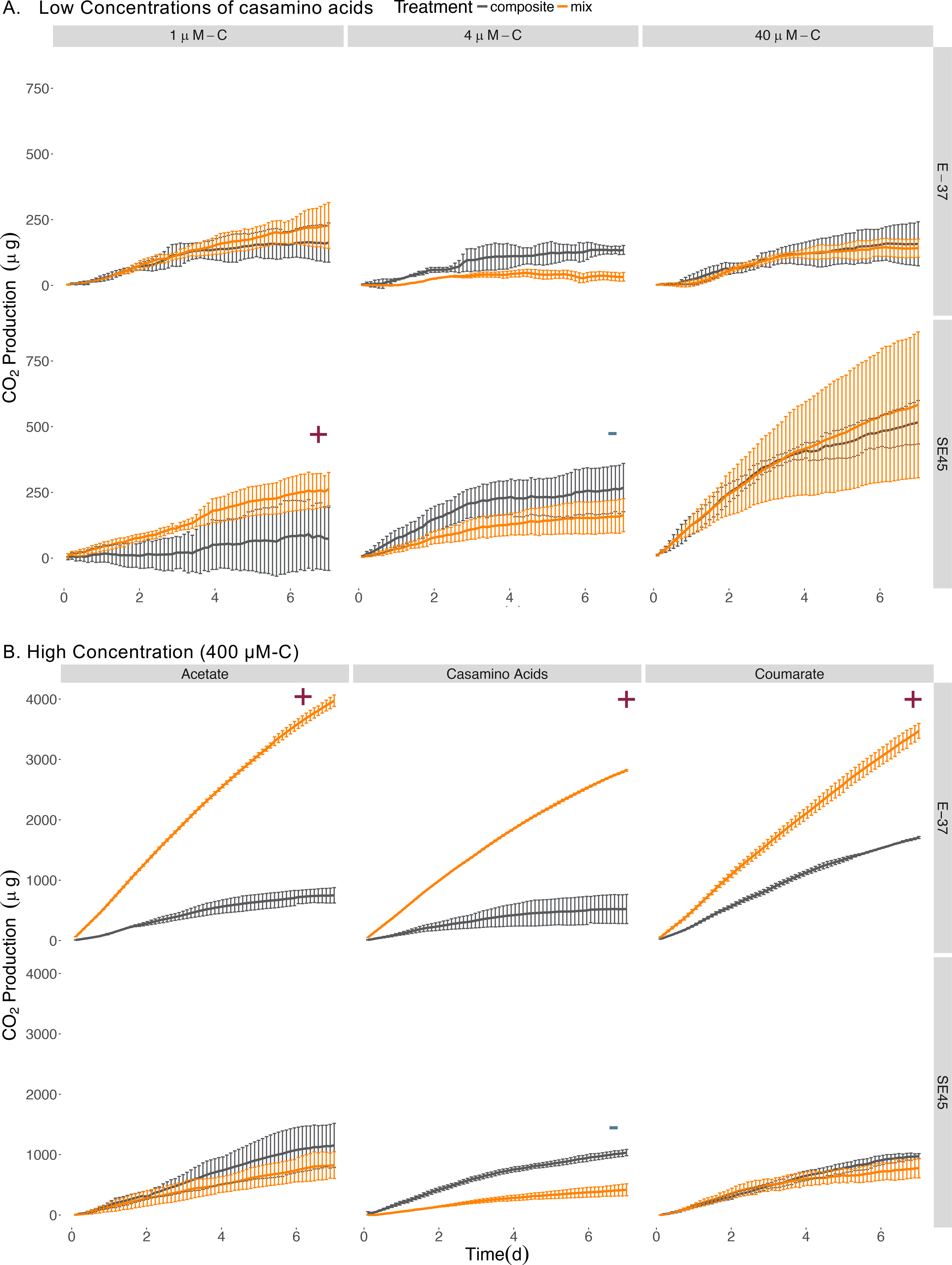
Cumulative CO_2_ production from SE45 and E-37 monocultures when provided (A) low concentrations of casamino acids and (B) high concentrations (400 μM-C) of acetate, casamino acids, and coumarate. Composite data (sum of LOM and NOM treatments) are shown in grey; mixed substrate data in orange. The average of the No C control was subtracted from all replicates. Points represent the mean (n=2-3); error bars represent one standard deviation from the mean. Red plus signs indicate a significant synergistic interactive effect (*p* < 0.05), blue minus signs indicate an antagonistic interactive effect (*p* < 0.05). The seeding densities for SE45 and E-37 were 3.05 × 10^4^ CFU/mL (±7.97 × 10^3^), 1.43 × 10^4^ CFU/mL (±4.71 × 10^3^), respectively. Final cumulative CO_2_ produced for all control treatments is shown in Table S4.

### Constructed community displays similar dynamics to single strains under mixed OM conditions

Given the differential response of individual strains to homogenous and mixed substrate conditions, we next tested a six-member constructed community, which included both strains, to assess interactive effects amongst community members with different metabolic capabilties. Similar to the single strain experiments, concentration and source of the LOM addition interacted significantly to determine viable cell densities at each time point in the 14-day experiment (Fig 1, Table S8). For each source of LOM, the viable cell densities produced at 400 μM-C were significantly greater than those at lower LOM concentrations (three-way ANOVA, n=5, p<0.001 for all time points). For mixed NOM + LOM substrate experiments, the community demonstrated a synergistic growth response, for at least one time point, to all LOM sources at 400 μM-C, and tryptone at 40 μM-C and 4 μM-C (Fig. S4, Table 4). Though in some treatments (e.g., 4 μM-C tryptone and acetate and 400 μM-C casamino acids) this was preceded by an initial antagonistic interactive response. Relative to the 400 uM concentrations, the community displayed a significant reduction in viable counts when supplied with each LOM source at 1 μM-C; the intervening concentrations showed varying responses. The six-member constructed community was best able to utilize tryptone for growth; the three other LOM types reached cell densities ~25% of tryptone-fed cultures (Fig. 1). By Day 14, communities showed ~60% decline in maximum viable cells on all of the substrates at 400 μM, except coumarate, for which there was 90% mortality (Fig 1).

**Table 4.** Probability values^a^ for constructed community interactive effects experimental data shown in Figure 1.

| **LOM Concentration<sup>b</sup>** |  |  |  |  |
| --- | --- | --- | --- | --- |
| **LOM Source** | **1 μM-C** | **4 μM-C** | **40 μM-C** | **400 μM-C** |
| **Acetate** | *Days 1, 4, 7*<br>*p < 0.05* | *Days 1, 10, 14*<br>*p < 0.05* | *Day 10 p < 0.01* | **Days 4, 10, 14**<br>**p < 0.05** |
| **Casamino Acids** | *Days 1, 2, 4, 7, 10, 14*<br>*p < 0.005* | No Significant Difference | *Days 1, 14*<br>*p < 0.001* | *Day 4 p < 0.01*<br>**Day 10 p < 0.01** |
| **Coumarate** | *Days 1, 2, 7, 10, 14*<br>*p < 0.001* | No Significant Difference | No Significant Difference | **Days 1, 7**<br>**p < 0.001** |
| **Tryptone** | *Days 1, 2, 7, 10, 14*<br>*p < 0.001* | *Days 1, 4 p < 0.05* | **Day 10**<br>**p < 0.001** | **Days 10, 14**<br>**p < 0.01** |

<sup>a</sup> For each day, a three-way ANOVA was performed to determine whether differences in cell densities were being driven by treatment, concentration or source of LOM. *p*-values are adjusted to correct for the false discovery rate using the Benjamini-Hochberg correction.
<sup>b</sup> Days listed are days in which there was a significant difference between the composite and mixed treatments. *p*-values for synergistic interactive effects are **bolded** and the corresponding table cell is shaded gray; *p*-values for antagonistic interactive effects are *italicized*.

**Table 5.** Probability values^a^ for respiration interactive effects experimental data shown in Figures 2 & 4.

| **LOM Concentration<sup>b</sup>** |  |  |  |  |
| --- | --- | --- | --- | --- |
| **LOM Source** | **1 μM-C** | **4 μM-C** | **40 μM-C** | **400 μM-C** |
| **Acetate** | Not Measured | Not Measured | Not Measured | SE45: $p < 0.30$<br>E-37: **$p < 0.001$**<br>Community: $p < 0.50$ |
| **Casamino Acids** | SE45: **$p < 0.05$**<br>E-37: $p < 0.33$<br>Community: $p < 0.6$ | SE45: $p < 0.25$<br>E-37: $p < 0.35$<br>Community: $p < 0.33$ | SE45: $p < 0.45$<br>E-37: $p < 0.95$<br>Community: *$p < 0.001$* | SE45: $p < 0.05$<br>E-37: **$p < 0.001$**<br>Community: $p < 0.60$ |
| **Coumarate** | Not Measured | Not Measured | Not Measured | SE45: $p < 0.55$<br>E-37: **$p < 0.001$**<br>Community: $p < 0.20$ |

<sup>a</sup> Due to the unbalanced nature of the respirometer experimental design, two three-way ANOVAs were used to analyze the differences between mix and composite in terms of final CO<sub>2</sub> accumulation. The two ANOVA models used tested whether the independent variables of inoculum, treatment, and either LOM source or concentration interacted to affect CO<sub>2</sub> accumulation. As the 400 μM-C Casamino Acids data were analyzed in both ANOVAs, the higher of the two resulting $p$ -value from the post hoc test were used to determine significance. $p$ -values are adjusted to correct for the false discovery rate using the Benjamini-Hochberg correction.
<sup>b</sup> $p$ -values for synergistic interactive effects are **bolded**; $p$ -values for antagonistic interactive effects are *italicized*.

No synergistic responses were observed in the constructed community when respiration was used as the measure of microbial activity. However, significant antagonistic responses were observed in the 40 μM-C casamino acids treatment (Table 5), as well as significantly lower CO_2_ production rates (1.5-fold) compared to the LOM treatment (three-way ANOVA, n=3, *p*<0.001, Tables S5-7), corroborating some of the antagonistic results from the viable count-based approach used in the first experiment (see Figs. 1 and 4). CO_2_ production rates were 1.3 and 1.5-fold higher in the mixed treatments than the LOM alone treatments for the low concentrations (1 and 4 μM-C) of casamino acids treatments (three-way ANOVA, n=3, *p*<0.001) and were statisitically indistinguishable at the highest LOM concentrations (Tables S5-7).

### LOM type drives microbial community composition

We next assessed the influence of concentration and source of LOM on community composition in the six-member culture. Species diversity decreased with increasing LOM concentration in both single and mixed OM substrate treatments (Fig. S5). At the highest LOM concentration, mesocosms were dominated by a single strain: either SE45 on coumarate or Y4I on the other three LOM types (Fig. 3). Treatment (mix or composite), LOM concentration, and LOM source interacted significantly to influence species diversity for all time points, with the exception of Day 2 where only LOM concentration and source interacted significantly (three-way ANOVA, n=5, *p*< 0.002 for all time points) (Table S9). LOM concentration, LOM source, and treatment interacted significantly to drive differences between communities throughout the course of the incubation (permutational MANOVA, *p*<0.05). Treatments using coumarate as LOM source resulted in a community distinct from the other sources of LOM at 400 μM-C (Fig. 3). Coumarate communities were characterized by increased abundances of SE45, comprising up to 84-90% of the community, compared to the other sources of LOM, where communities were dominated by strain Y4I (up to 85-98% of the community) (Fig. 3). Mixed LOM + NOM treatments had increased viable cell abundances for E-37 and SE45 compared to LOM alone treatments and both of these strains have increased viable cell densities in the NOM alone treatments compared to No C (Figs. 1 and 3). The community composition within the respirometer experiments generally mirrored that of the viable counts experiment in which Y4I was the most abundant member of the community for all sources of LOM, with the exception of the coumarate treatments (Fig. S6). The most notable difference between the community composition in the viable count vs respirometer experiments is that Y4I maintained higher relative abundances at lower concentrations of LOM than in the viable counts (Fig. S6).

**Figure 3.**
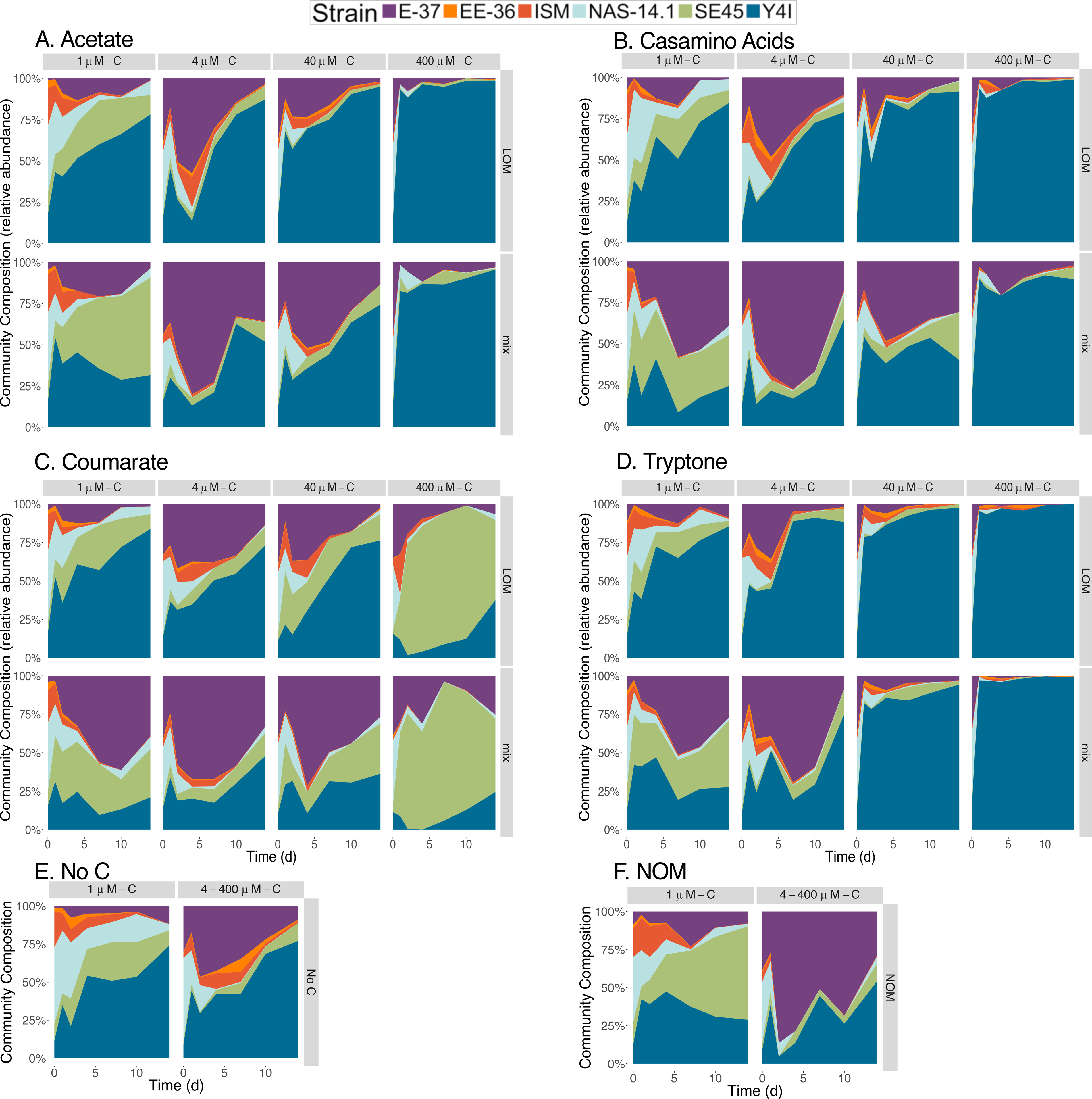
Community composition of the six-member constructed community in response to treatments with varying concentrations of (A) acetate (B) casamino acids, (C) coumarate and (D) tryptone. Community composition is displayed in relative abundance and individual strains are color- coded according to the key. Each LOM alone treatment is displayed above its corresponding mix treatment. No carbon and NOM alone controls are shown in panels (E) and (F), respectively. The paired NOM and No C treatment community compositions for the 1 uM LOM concentrations are shown on the left and those for the for the 4, 40 and 400 uM are shown on the right. Seeding density for the constructed communities was 7.01 × 103 CFU/mL (±2.6 × 103).

**Figure 4.**
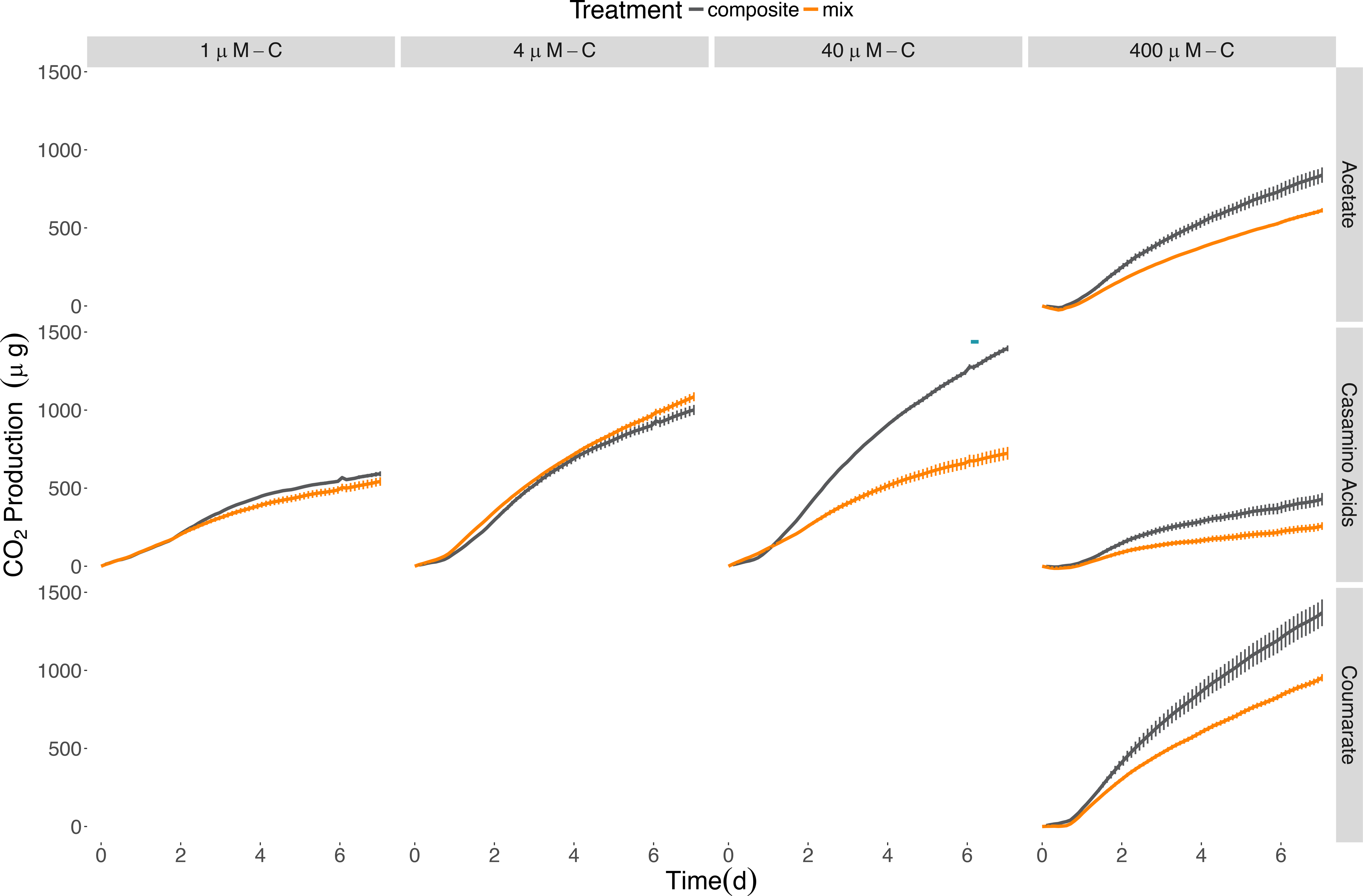
Cumulative CO_2_ production for the six-member constructed community provided different sources and concentrations of LOM. Composite data (sum of LOM and NOM) are shown in grey; mixed substrate data in orange. The average of the No C control was subtracted from all replicates. Points represent the mean (n=2-3) while error bars represent one standard deviation from the mean. Red plus signs indicate a significant synergistic interactive effect (*p* < 0.05) and blue minus signs indicate an antagonistic interactive effect (*p* < 0.05). The seeding density was 5.13 × 10^3^ CFU/mL (± 3.73 × 10^3^). Final cumulative CO_2_ produced for all control treatments is shown in Table S4.

## Discussion

Traditionally, geochemical models designate organic matter as consisting of multiple, independent pools of compounds, each of which is degraded by microorganisms at different rates (Arndt *et al.* 2013; Hansell 2013). However, recent findings in microbial physiology suggest that organic compounds can interact within microbial metabolisms in unpredictable ways (Gulvik and Buchan 2013; Gontikaki *et al.* 2015; Ward *et al.* 2016). The degree to which OM interactivity in general, and the priming effect (PE) specifically, are quantitatively important in aquatic ecosystems is an area of current study and debate. Field and lab studies have shown that, depending on the precise circumstance, labile organic matter can speed, slow, or have no effect on the oxidation of recalcitrant organic matter in aquatic environments (e.g., Bengtsson *et al.* 2014; Gontikaki *et al.* 2013; Bianchi *et al.* 2015; Catalán *et al.* 2015; Steen, Quigley and Buchan 2016). Given these inconsistencies in the literature, we set out to perform controlled laboratory experiments to characterize interactive effects of distinct OM pools on microbial metabolism.

Coastal salt marsh microbial communities are subject to periodic pulses of OM from multiple sources and are inherently complex (e.g. Moran *et al.* 2007; Medeiros *et al.* 2017). Thus, the use of environmentally relevant and culturable representatives from these communities provides a tractable system for obtaining foundational knowledge of the underlying mechanisms of interactive effects on microbial processing of OM. Here, we used cultured representatives from a lineage of coastal marine bacteria that are known to dominate and be metabolically active in coastal estuaries (Buchan, González and Moran 2005; Bakenhus *et al.* 2017). These bacteria were provided a natural and environmentally relevant source of recalcitrant organic matter, natural organic matter (NOM) derived from a river feeding Southeastern US coastal estuaries. We assessed the microbial metabolic response to mixtures of labile and recalcitrant OM in two ways: by measuring viable cell abundance and by measuring CO_2_ production. These experiments revealed the importance of labile substrate concentration and chemical composition in dictating the growth dynamics of representative marine bacteria in the presence of natural organic matter. We quantified species-specific responses to mixed substrate regimes, documented the transient nature of these responses and demonstrated microbial community composition shifts in response to interactive effects in relevant mixed carbon conditions.

### Interactive OM effects are often transient

Our data with cultured bacteria demonstrate evidence of the transience of interactivity in OM degradation. These interactivities occurred on timeframes consistent with what has been reported previously in the PE literature for both individual microbial isolates and communities (1-7 days e.g., D’Errico *et al.* 2013; Bianchi *et al.* 2015; Steen, Quigley and Buchan 2016). Synergistic interactive effects of the labile and recalicitrant C sources on microbial growth were detectable either within the first few days of our incubations and/or as the microbial populations started to decline towards the end of the 14-day experimental period. With few exceptions, when synergistic interactions occurred, they did not persist beyond a two to four-day timeframe. With some treatments, an additional, temporally distinct and positive synergistic interaction was observed 10-14 days into the experiment. Antagonistic interactive effects were also observed: for SE45 in half of the treatments conditions and ~40% of the treatments for E-37 and the constructed community. These treatments were almost always at the lower concentrations of LOM, with the exception of E-37 when provided 400 μM-C. While most of the antagonistic effects are transient, SE45 produces long-lasting antagonistic responses when provided NOM with 4 μM-C casamino acids and coumarate. E-37 demonstrated antagonistic effects lasting almost the entirety of the incubation when provided NOM along with 4 μM-C coumarate, and 4 or 400 μM-C tryptone. The constructed community displayed long antagnostic effects when provided 1 μM-C casamino acids, coumarate, and tryptone. Mix conditions conditions containing acetate did not elicit long-lasting antagonistic effects. Instead, acetate appears to stimulate synergistic effects of the greatest duration in each inocula. This may indicate that, for Roseobacters, chemically simple labile substrates that feed directly into central metabolism are more likely to stimulate a synergistic effect at high concentrations, while more complex labile substrates may result in an antagonistic response at low concentrations.

While viable cell densities in treatments with NOM alone stayed relatively consistent throughout the incubation, most of the LOM only treatments exhibited severe declines in viable cell densities following an initial increase in cell growth. Those declines drove down the composite values used for comparisons with mix treatments to quantitatively assess interactive effects. The degree of viable cell demise tended to increased with increasing substrate concentration. Given the time scales of the experiments performed here, it is unlikely that substantial numbers of cells died as a result of lack of organic carbon or nutrients (Novitsky and Morita 1978). Furthermore, the medium was well-buffered to prevent dramatic changes in pH. However, it is plausible that the catabolism of a given LOM and/or NOM results in the production of toxic metabolic by-product, giving rise to decreased viability. It has been previously shown that microbial conversion of simple carbon substrates can result in the production and release of a diversity of compounds (Lechtenfeld *et al.* 2015). Alternatively, prophage induction could have contributed to mortality. Three of the six strains, E-37, SE45, and Y4I, are predicated to encode prophage (Table 2), though to our knowledge induction of these prophages has not yet been demonstrated for any of these putative lysogens. Nonetheless, recent evidence suggests a correlation between bacterial productivity and lysogenic to lytic conversion in natural systems (Brum *et al.* 2016). Indeed, these two ideas are not mutually exclusive as increased bacterial metabolism could lead to enhanced production of toxic metabolic by-products that could, in turn, induce a global stress response in bacterial strains, initiating a lysogenic-lytic conversion (Feiner *et al.* 2015). Finally, growth substrate has been shown to influence prophage induction, indicating host metabolic state can have a direct influence on the lysogenic-lytic decision (e.g. Howes 1965; Czyz *et al.* 2001).

Viable cell densities did not decline in the mixed treatments as precipitously as those in the LOM alone treatments. The apparent stabilizing effect seen in the mixed OM treatments compared to the composites may arise from the ability of the bacteria to access additional components of NOM, enabled by LOM catabolism, a mechanism posited by Guenet and colleagues (2010). Furthermore, in some instances, the cell densities in the mixed treatments begin to rebound towards the later stages of the experiment. The mixed carbon regime provided by the combination of LOM and NOM may yield conditions favorable for microbial adaptations, such as the proliferation of growth advantage in stationary phase (GASP) mutants (Zinser and Kolter 1999), which could utilize previously unavailable components of the NOM, or possibly tolerate toxic compounds released by actively growing cells earlier in the incubation. Additional experiments are needed to specifically address the contribution of microbial adaptation, acclimation, prophage induction and metabolite toxicity to the observed trends.

Some inconsistencies between interactive effects in the viable counts and respiration data were observed and not completely unexpected. These discrepancies may indicate altered growth efficiencies under different substrate regimes. Alternatively, cultures had to remain static in the respirometer and it is plausible that biofilms developed under these conditions, though they were not visible to the naked eye. Roseobacters are prolific in natural marine biofilms (Dang and Lovell 2000; Dang *et al.* 2008) and all six of these strains have been previously demonstrated to form biofilms when grown on complex media (Slightom & Buchan, 2009). The physiological status of bacteria growing in biofilms is different from those grown planktonically due to alterations in gene expression that can lead to changes in cell surface chemistry, physiology and behavior (Costerton *et al.* 1995). Thus, surface associated growth could influence microbial catabolism under mixed substrate regimes. Additional studies are needed to tease apart the contributions of these factors and we caution against making direct comparisons between the experiments that relied on viable counts (shown in Figs 1 & 3) and those that monitored respiration (shown in Figs 2 & 4).

### Interactive effects are species-specific

While there is overlap between the mixed carbon substrate conditions that stimulate or repress growth of SE45, E-37 and the constructed community, each inoculum experienced OM interactivity under a unique set of conditions. For example, SE45 demonstrated a synergistic response to mixtures of NOM with 400 μM-C tryptone, a treatment in which E-37 responded antagonistically. The differential ability of SE45 and E-37 to undergo synergistic interactive effects through the addition of tryptone suggests that the expression and/or activity of extracellular enzymes could be an important factor in the onset of interactive effects. While monocultures of E-37 ultimately reach similar viable cell densities as all other members of the community, E-37 displays a considerably longer lag phase relative to the other strains when grown on 2 mM-C tryptone as a monoculture (Fig. S7). The delayed growth on tryptone may prevent E-37 from exhibiting a synergistic response with tryptone plus NOM when grown by itself. However, E-37 outperforms SE45 in the constructed community when provided low concentration of tryptone, which as discussed below, is indicative of synergistic interactions between community members. Both E-37 and SE45 undergo a synergistic interactive effect in the 40 μM-C acetate mixed treatment. However, the community undergoes an antagonistic interacive effect under the same conditions. It is plausible that competition with Y4I, the overwhelming dominant member of the community at high LOM concentrations of tryptone, casamino acids and acetate, contributes to the antagonistic effect seen in the constructed community treatments by preventing either E-37 or SE45 from performing necessary metabolic processes required for a synergistic response.

In agreement with our earlier report that a natural estuarine microbial community underwent a significant positive interactive effect with the addition of a globular protein (bovine serum albumin, provided at 500 μM) (Steen, Quigley and Buchan 2016), the constructed community analyzed in this study displayed synergistic interactive effect in the presence of tryptone, an assortment of peptides, at 400 μM. However, timing of a response differed: it was delayed in the the constructed community with tryptone (occurring during the second week of incubation) compared to an immediate priming response by the natural community provided complex protein. While there are many factors that could contribute to this apparent temporal disconnect, the relatively low strain diversity of the constructed community may be a key driver. By day 1, the constructed community was dominanted by a single strain: Y4I comprised 98% of the community in this treatment. Y4I belongs to the genus *Phaeobacter*, members of which were recently shown to bloom in the presence of Arctic riverine, dissolved organic matter (Sipler *et al.* 2017). Additionally, we earlier observed that acetate (at 500 μM-C) repressed the ability of a estuarine microbial community to degrade phytoplankton necromass (Steen, Quigley and Buchan 2016). However, in the current experiments all bacterial inocula demonstrated a synergistic growth response to the addition of acetate, at the highest concentration (400 μM-C). Collectively, these findings demonstrate that the substrate conditions that result in OM interactive effects are species-specific and thus dictated by the composition and metabolic potential of a community.

### Carbon sources shape the composition and diversity of the constructed community

While scant information exists on how interactive effects influence community composition, studies that indicate riverine DOM structures the composition of microbial communities along the river-estuary continuum provide a useful comparative framework (Langenheder *et al.* 2004; Blanchet *et al.* 2017). One report using an estuarine community incubated with riverine DOM and casamino acids saw no evidence for interactive effects and only minor alterations in microbial community composition (Blanchet *et al.* 2017). In contrast, we observed that the diversity of our constructed microbial community was influenced significantly both by the carbon sources present (e.g. LOM, NOM, or mixtures of the two) and the concentrations and sources of the LOM (Table S9). E-37 has been previously shown to simultaneously catabolize aromatic compounds via two different ring cleaving pathways, the benzoyl Co-A and protocatechuate pathways, and derive a beneficial effect when grown on a mixture of carbon substrates compared to either substrate presented alone (Gulvik and Buchan 2013). The metabolic synergy between these two aromatic carbon catabolism pathways may also be a mechanism for OM interactivities that has been previously overlooked.

Our studies reveal that structure of the constructed communities may often be determined by the concentration of LOM provided, regardless of chemical form. With the exception of the highest coumarate concentration treatment, a general trend emerged: as the concentration of LOM increases, the diversity within the constructed community decreases. This stands in contrast to some prior studies in which increasing amounts of autochthonous carbon resulted in increased degradation of allochthounous carbon, with little to no effect on bacterial community composition (Attermeyer *et al.* 2014). This decrease in diversity was most pronounced in the highest LOM additions (400 μM-C), where a single strain (Y4I) dominated all, but the coumarate, treatments. The shorter lag phase and faster growth rate of Y4I relative to other members of the community when grown on labile substrates may have allowed Y4I to gain an early foothold in the community. This possibility is supported by the fact that the numerical dominace of Y4I began as early as day 1 in the incubations, after which it either increased in terms of relative abundance or maintained its numerical dominance in the community (Fig. 3). The stark contrast in community composition between those cultures provided coumarate compared to the other LOM types is likely due to the unique ability of SE45 and E-37 to utilize coumarate as a carbon source. For the coumarate treatments, SE45, and to a lesser extent E-37, become the most numerically abundant organisms. Given that these strains are both ligninolytic they are likely better tuned to access the aromatic carbon moieties characteristic of NOM (Gonzalez *et al.* 1997; Frank 2016).

Cooperation and competition may be important ecological processes influencing the outcome of interactive effects (Fontaine, Mariotti and Abbadie 2003). Our data indicate both cooperation and competition under different conditions in the constructed community experiments. For example, SE45 reached higher cell densities in the constructed community in the presence of both NOM and 400 μM-C coumarate compared to its growth on these substrates in monoculture. Additionally, E-37 growth is enhanced in the constructed community when provided low concentrations of tryptone compared to monocultures. This strain may gain an advantage from other members of the community that produce extracellular peptidases, liberating free amino acids that E-37 is, in turn, more competitive at transporting and catabolizing. While many bacteria can transport the lower molecular weight fraction of tryptone directly into cells via oligopeptide permeases (Garault *et al.* 2002), up to 10% of tryptone is between 2-5 kDa (BD Biosciences 2006) and requires initial cleavage by extracellular peptidases. Extracellular enzymes are generally considered “public goods” because they may provide benefit to the community, while being costly for individuals to produce. Individuals within a community who take advantage of public goods without producing them are termed cheaters and cheating has been shown to increase in frequency in well-mixed systems with high diffusion rates (Allison *et al.* 2014), such as the culture conditions employed in this study.

In coastal marshes, the dissolved organic carbon pool is highly heterogenous in both structure and distribution. Similarly, the microbial communities in these systems display a high degree of genetic and functional diversity and are patchy in both their abundances and activity. Deciphering the complex chemical and biological interactions that unlie the mineralization of organic carbon in these systems is a daunting challenge. Yet, a detailed understanding of the nature and sources of LOM to estuaries, as well as the molecular mechanisms driving interactive effects, will be necessary to understand the controls on microbial oxidation of terrestrial organic carbon in estuaries. In order to elucidate microbe-multi-substrate interactivities, controlled laboratory experiments employing relatively low chemical and biological complexity provide an important foundation on which to build.

## Acknowledgments

We thank Terry Hazen for the use of the respirometer and Melanie Mayes and Hannah Woo for training on the instrument.

## Funding

This research was supported by a grant from the National Science Foundation [OCE-1357242] awarded to AB and ADS.

