## Supplemental Tables for "Characterization of the interactive effects of labile and recalcitrant organic matter on microbial growth and metabolism"

#### Supplemental Tables (Quigley et al)

**Supplemental Table S1. ANOVA Tables for each three-way ANOVA performed for SE45 cell density by day.**

| Day | ANOVA Table | | | | | |
|---|---|---|---|---|---|---|
| 1 | | Df | Sum Sq | Mean Sq | F value | Pr(>F) |
| | Treatment | 1 | 0.14 | 0.145 | 3.866 | 0.05143 |
| | Concentration | 3 | 46.41 | 15.470 | 413.565 | < 2e-16 |
| | Carbon | 3 | 0.90 | 0.300 | 8.011 | 6.19e-05 |
| | Treatment:Concentration | 3 | 1.75 | 0.582 | 15.556 | 1.10e-08 |
| | Treatment:Carbon | 3 | 0.46 | 0.154 | 4.110 | 0.00804 |
| | Concentration:Carbon | 9 | 3.49 | 0.388 | 10.372 | 6.39e-12 |
| | Treatment:Concentration:Carbon | 9 | 2.21 | 0.246 | 6.566 | 1.14e-07 |
| | Residuals | 128 | 4.79 | 0.037 | | |
| 2 | | Df | Sum Sq | Mean Sq | F value | Pr(>F) |
| | Treatment | 1 | 0.40 | 0.397 | 14.620 | 0.000211 |
| | Concentration | 3 | 51.95 | 17.318 | 638.473 | < 2e-16 |
| | Carbon | 3 | 2.15 | 0.715 | 26.378 | 3.75e-13 |
| | Treatment:Concentration | 3 | 0.96 | 0.320 | 11.812 | 7.92e-07 |
| | Treatment:Carbon | 3 | 0.03 | 0.010 | 0.351 | 0.788349 |
| | Concentration:Carbon | 9 | 3.22 | 0.358 | 13.203 | 1.90e-14 |
| | Treatment:Concentration:Carbon | 9 | 0.31 | 0.034 | 1.271 | 0.259552 |
| | Residuals | 119 | 3.23 | 0.027 | | |
| 4 | | Df | Sum Sq | Mean Sq | F value | Pr(>F) |
| | Treatment | 1 | 0.67 | 0.666 | 28.833 | 3.70e-07 |
| | Concentration | 3 | 32.28 | 10.760 | 465.940 | < 2e-16 |
| | Carbon | 3 | 0.75 | 0.250 | 10.836 | 2.23e-06 |
| | Treatment:Concentration | 3 | 0.59 | 0.196 | 8.500 | 3.50e-05 |
| | Treatment:Carbon | 3 | 0.47 | 0.156 | 6.738 | 0.000299 |
| | Concentration:Carbon | 9 | 5.08 | 0.564 | 24.423 | < 2e-16 |
| | Treatment:Concentration:Carbon | 9 | 0.70 | 0.078 | 3.361 | 0.001020 |
| | Residuals | 125 | 2.89 | 0.023 | | |
| 7 | | Df | Sum Sq | Mean Sq | F value | Pr(>F) |
| | Treatment | 1 | 0.003 | 0.003 | 0.100 | 0.752828 |
| | Concentration | 3 | 18.786 | 6.262 | 210.239 | < 2e-16 |
| | Carbon | 3 | 0.501 | 0.167 | 5.606 | 0.001219 |
| | Treatment:Concentration | 3 | 0.387 | 0.129 | 4.332 | 0.006084 |
| | Treatment:Carbon | 3 | 1.031 | 0.344 | 11.539 | 9.82e-07 |
| | Concentration:Carbon | 9 | 3.736 | 0.415 | 13.939 | 2.23e-15 |
| | Treatment:Concentration:Carbon | 9 | 1.001 | 0.111 | 3.734 | 0.000347 |
| | Residuals | 126 | 3.753 | 0.030 | | |
| 10 | | Df | Sum Sq | Mean Sq | F value | Pr(>F) |
| | Treatment | 1 | 0.020 | 0.020 | 0.643 | 0.424108 |
| | Concentration | 3 | 15.930 | 5.310 | 172.504 | < 2e-16 |
| | Carbon | 3 | 0.643 | 0.214 | 6.959 | 0.000232 |
| | Treatment:Concentration | 3 | 1.206 | 0.402 | 13.060 | 1.88e-07 |
| | Treatment:Carbon | 3 | 0.399 | 0.133 | 4.321 | 0.006221 |
| | Concentration:Carbon | 9 | 1.131 | 0.126 | 4.083 | 0.000133 |
| | Treatment:Concentration:Carbon | 9 | 0.843 | 0.094 | 3.042 | 0.002571 |
| | Residuals | 122 | 3.755 | 0.031 | | |
| 14 | | Df | Sum Sq | Mean Sq | F value | Pr(>F) |
| | Treatment | 1 | 0.217 | 0.2170 | 4.739 | 0.031419 |
| | Concentration | 3 | 9.323 | 3.1078 | 67.872 | < 2e-16 |
| | Carbon | 3 | 2.528 | 0.8427 | 18.405 | 6.48e-10 |
| | Treatment:Concentration | 3 | 1.183 | 0.3942 | 8.609 | 3.14e-05 |
| | Treatment:Carbon | 3 | 0.192 | 0.0638 | 1.394 | 0.247777 |
| | Concentration:Carbon | 9 | 1.714 | 0.1904 | 4.158 | 0.000107 |
| | Treatment:Concentration:Carbon | 9 | 0.420 | 0.0467 | 1.020 | 0.428257 |
| | Residuals | 122 | 5.586 | 0.0458 | | |

**Supplemental Table S2. ANOVA Tables for each three-way ANOVA performed for E-37 cell density by day.**

| Day | ANOVA table | | | | | |
|---|---|---|---|---|---|---|
| 1 | | Df | Sum Sq | Mean Sq | F value | Pr(>F) |
| | Treatment | 1 | 0.004 | 0.004 | 0.269 | 0.60469 |
| | Concentration | 3 | 28.549 | 9.516 | 649.373 | < 2e-16 |
| | Carbon | 3 | 1.715 | 0.572 | 38.999 | < 2e-16 |
| | Treatment:Concentration | 3 | 0.231 | 0.077 | 5.257 | 0.00192 |
| | Treatment:Carbon | 3 | 0.620 | 0.207 | 14.102 | 6.09e-08 |
| | Concentration:Carbon | 9 | 3.846 | 0.427 | 29.159 | < 2e-16 |
| | Treatment:Concentration:Carbon | 9 | 1.185 | 0.132 | 8.983 | 2.67e-10 |
| | Residuals | 121 | 1.773 | 0.015 | | |
| 2 | | Df | Sum Sq | Mean Sq | F value | Pr(>F) |
| | Treatment | 1 | 0.017 | 0.017 | 0.466 | 0.49587 |
| | Concentration | 3 | 15.543 | 5.181 | 139.414 | < 2e-16 |
| | Carbon | 3 | 4.327 | 1.442 | 38.814 | < 2e-16 |
| | Treatment:Concentration | 3 | 0.419 | 0.140 | 3.755 | 0.01266 |
| | Treatment:Carbon | 3 | 0.633 | 0.211 | 5.682 | 0.00111 |
| | Concentration:Carbon | 9 | 2.536 | 0.282 | 7.581 | 7.93e-09 |
| | Treatment:Concentration:Carbon | 9 | 0.412 | 0.046 | 1.233 | 0.28075 |
| | Residuals | 126 | 4.682 | 0.037 | | |
| 4 | | Df | Sum Sq | Mean Sq | F value | Pr(>F) |
| | Treatment | 1 | 0.190 | 0.190 | 2.547 | 0.11314 |
| | Concentration | 3 | 11.964 | 3.988 | 53.593 | < 2e-16 |
| | Carbon | 3 | 5.806 | 1.935 | 26.010 | 4.70e-13 |
| | Treatment:Concentration | 3 | 0.263 | 0.088 | 1.177 | 0.32142 |
| | Treatment:Carbon | 3 | 1.062 | 0.354 | 4.758 | 0.00359 |
| | Concentration:Carbon | 9 | 3.548 | 0.394 | 5.297 | 4.38e-06 |
| | Treatment:Concentration:Carbon | 9 | 2.056 | 0.228 | 3.070 | 0.00238 |
| | Residuals | 121 | 9.004 | 0.074 | | |
| 7 | | Df | Sum Sq | Mean Sq | F value | Pr(>F) |
| | Treatment | 1 | 2.255 | 2.255 | 57.171 | 7.73e-12 |
| | Concentration | 3 | 0.294 | 0.098 | 2.486 | 0.0637 |
| | Carbon | 3 | 11.767 | 3.922 | 99.457 | < 2e-16 |
| | Treatment:Concentration | 3 | 0.142 | 0.047 | 1.200 | 0.3126 |
| | Treatment:Carbon | 3 | 1.493 | 0.498 | 12.618 | 2.97e-07 |
| | Concentration:Carbon | 9 | 5.829 | 0.648 | 16.422 | < 2e-16 |
| | Treatment:Concentration:Carbon | 9 | 2.880 | 0.320 | 8.113 | 2.13e-09 |
| | Residuals | 124 | 4.890 | 0.039 | | |
| 10 | | Df | Sum Sq | Mean Sq | F value | Pr(>F) |
| | Treatment | 1 | 0.006 | 0.0056 | 0.101 | 0.7513 |
| | Concentration | 3 | 5.987 | 1.9957 | 35.790 | < 2e-16 |
| | Carbon | 3 | 9.430 | 3.1432 | 56.370 | < 2e-16 |
| | Treatment:Concentration | 3 | 1.624 | 0.5414 | 9.709 | 8.91e-06 |
| | Treatment:Carbon | 3 | 2.209 | 0.7364 | 13.206 | 1.71e-07 |
| | Concentration:Carbon | 9 | 4.080 | 0.4534 | 8.131 | 2.56e-09 |
| | Treatment:Concentration:Carbon | 9 | 1.112 | 0.1235 | 2.216 | 0.0255 |
| | Residuals | 118 | 6.580 | 0.0558 | | |
| 14 | | Df | Sum Sq | Mean Sq | F value | Pr(>F) |
| | Treatment | 1 | 0.110 | 0.110 | 2.995 | 0.0861 |
| | Concentration | 3 | 20.854 | 6.951 | 189.836 | < 2e-16 |
| | Carbon | 3 | 3.376 | 1.125 | 30.733 | 7.69e-15 |
| | Treatment:Concentration | 3 | 0.212 | 0.071 | 1.926 | 0.1290 |
| | Treatment:Carbon | 3 | 1.602 | 0.534 | 14.582 | 3.62e-08 |
| | Concentration:Carbon | 9 | 1.500 | 0.167 | 4.551 | 3.55e-05 |
| | Treatment:Concentration:Carbon | 9 | 0.724 | 0.080 | 2.197 | 0.0267 |
| | Residuals | 121 | 4.431 | 0.037 | | |

**Supplemental Table S3. Mean viable counts[a] for Day 7 of respirometer experiments for all treatments.**

| | SE45 Respirometer[b] | E-37 Respirometer[c] | Community Respirometer[d] |
|---|---|---|---|
| No C | $1.34 \times 10^{5} \pm 1.38 \times 10^{5}$ | $2.28 \times 10^{4} \pm 6.55 \times 10^{3}$ | $6.63 \times 10^{5} \pm 2.57 \times 10^{5}$ |
| NOM | $2.79 \times 10^{5} \pm 1.48 \times 10^{5}$ | $2.17 \times 10^{5} \pm 6.41 \times 10^{4}$ | $2.57 \times 10^{6} \pm 2.57 \times 10^{5}$ |
| 1 µM-C Casamino LOM | $3.97 \times 10^{5} \pm 5.42 \times 10^{4}$ | $5.73 \times 10^{4} \pm 6.66 \times 10^{3}$ | $6.37 \times 10^{5} \pm 7.02 \times 10^{4}$ |
| 1 µM-C Casamino mix | $4.30 \times 10^{5} \pm 2.17 \times 10^{4}$ | $2.56 \times 10^{5} \pm 5.28 \times 10^{4}$ | $1.51 \times 10^{6} \pm 1.11 \times 10^{5}$ |
| 4 µM-C Casamino LOM | $3.53 \times 10^{4} \pm 1.06 \times 10^{4}$ | $3.33 \times 10^{5} \pm 4.07 \times 10^{5}$ | $8.03 \times 10^{6} \pm 1.91 \times 10^{6}$ |
| 4 µM-C Casamino mix | $1.78 \times 10^{5} \pm 9.05 \times 10^{4}$ | $3.30 \times 10^{5} \pm 4.00 \times 10^{4}$ | $2.26 \times 10^{7} \pm 2.88 \times 10^{6}$ |
| 40 µM-C Casamino LOM | $1.49 \times 10^{5} \pm 5.82 \times 10^{4}$ | $3.10 \times 10^{5} \pm 3.91 \times 10^{5}$ | $2.15 \times 10^{7} \pm 2.40 \times 10^{6}$ |
| 40 µM-C Casamino mix | $4.71 \times 10^{5} \pm 1.21 \times 10^{5}$ | $3.40 \times 10^{5} \pm 7.55 \times 10^{4}$ | $3.27 \times 10^{6} \pm 4.62 \times 10^{5}$ |
| 400 µM-C Casamino LOM | $4.40 \times 10^{5} \pm 1.61 \times 10^{5}$ | $2.93 \times 10^{5} \pm 4.73 \times 10^{4}$ | $1.27 \times 10^{7} \pm 2.40 \times 10^{6}$ |
| 400 µM-C Casamino mix | $1.58 \times 10^{6} \pm 8.13 \times 10^{5}$ | $7.53 \times 10^{5} \pm 3.21 \times 10^{4}$ | $2.09 \times 10^{7} \pm 2.12 \times 10^{6}$ |
| 400 µM-C Acetate LOM | $1.07 \times 10^{7} \pm 3.96 \times 10^{5}$ | $2.27 \times 10^{5} \pm 7.09 \times 10^{4}$ | $9.87 \times 10^{6} \pm 9.29 \times 10^{5}$ |
| 400 µM-C Acetate mix | $8.31 \times 10^{6} \pm 1.08 \times 10^{6}$ | $8.97 \times 10^{5} \pm 7.64 \times 10^{4}$ | $1.74 \times 10^{7} \pm 2.29 \times 10^{6}$ |
| 400 µM-C Coumarate LOM | $2.55 \times 10^{6} \pm 4.72 \times 10^{5}$ | $5.57 \times 10^{5} \pm 5.86 \times 10^{4}$ | $1.45 \times 10^{6} \pm 2.12 \times 10^{5}$ |
| 400 µM-C Coumarate mix | $3.96 \times 10^{6} \pm 6.41 \times 10^{5}$ | $7.47 \times 10^{5} \pm 6.11 \times 10^{4}$ | $3.18 \times 10^{6} \pm 1.60 \times 10^{6}$ |

[a]The mean and one standard deviation are reported. No C and NOM CFU/mL was calculated from n=6, while the CFU/mL for the mix and LOM treatments was calculated from n=3.
[b]The seeding density for the SE45 respirometer experiments was $3.05 \times 10^{4}$ CFU/mL ($\pm 7.97 \times 10^{3}$).
[d]The seeding density for the E-37 respirometer experiments was $1.43 \times 10^{4}$ CFU/mL ($\pm 4.71 \times 10^{3}$).
[f]The seeding density for the community respirometer experiments was $5.13 \times 10^{3}$ CFU/mL ($\pm 3.73 \times 10^{3}$).

**Supplemental Table S4. Average amount of $CO_2$ (μg) respired by the final time (Day 7) point for each treatment.**

| Treatment | composite | LOM | NOM[a] | mix |
|---|---|---|---|---|
| SE45<br>400 μM-C Acetate | 1153.63±364.83 | 963.72±415.24 | 189.91±51.34 | 826.51±221.18 |
| SE45<br>1 μM-C Casamino Acids | 73.39±120.38 | -116.52±125.78 | 189.91±51.34 | 262.94±62.97 |
| SE45<br>4 μM-C Casamino Acids | 300.52±109.89 | 63.31±31.57 | 158.66±168.97 | 190.11±84.88 |
| SE45<br>40 μM-C Casamino Acids | 591.74±130.24 | 354.53±227.55 | 158.66±168.97 | 695.02±358.06 |
| SE45<br>400 μM-C Casamino Acids | 1283.08±76.02 | 1045.87±53.91 | 158.66±168.97 | 550.02±155.51 |
| SE45<br>400 μM-C Coumarate | 974.44±42.41 | 784.53±61.92 | 189.91±51.34 | 777.09±161.76 |
| E-37<br>400 μM-C Acetate | 750.16±127.13 | 426.02±147.20 | 324.12±175.10 | 3972.30±94.83 |
| E-37<br>1 μM-C Casamino Acids | 162.75±74.71 | 8.50±5.23 | 154.24±78.24 | 227.88±86.90 |
| E-37<br>4 μM-C Casamino Acids | 134.84±17.07 | -19.41±72.83 | 154.24±78.24 | 32.69±16.46 |
| E-37<br>40 μM-C Casamino Acids | 158.12±83.59 | 3.882±39.58 | 154.24±78.24 | 141.60±34.70 |
| E-37<br>400 μM-C Casamino Acids | 522.59±241.11 | 198.45±134.40 | 324.12±175.10 | 2817.89±13.65 |
| E-37<br>400 μM-C Coumarate | 1706.55±17.47 | 1289.16±78.09 | 324.12±175.10 | 3473.37+122.56 |
| Constructed Community<br>400 μM-C Acetate | 838.13±49.20 | 525.05±15.70 | 313.08±33.90 | 612.50 ±14.20 |
| Constructed Community<br>1 μM-C Casamino Acids | 591.67±16.17 | 293.57±20.93 | 298.10±4.80 | 542.65±27.33 |
| Constructed Community<br>4 μM-C Casamino Acids | 1000.83±32.32 | 702.73±28.38 | 298.10±4.80 | 1085.36±28.65 |
| Constructed Community<br>40 μM-C Casamino Acids | 1395.32±18.69 | 1097.22±23.49 | 298.10±4.80 | 723.12±40.75 |
| Constructed Community<br>400 μM-C Casamino Acids | 429.31±39.80 | 116.23±6.91 | 313.08±33.90 | 257.37±24.35 |
| Constructed Community<br>400 μM-C Coumarate | 1368.75±85.47 | 1055.67±73.57 | 313.08±33.90 | 952.79±23.31 |

[a] Samples were processed in batch. Variation in NOM alone values represents run-to-run variation as these controls were included in every batch run.

**Supplemental Table S5. ANOVA Tables for the three-way ANOVAs used to analyze the respiration data.**

| Test[a] | ANOVA Table | Df | Sum Sq | Mean Sq | F value | Pr(>F) |
|---|---|---|---|---|---|---|
| Accumulation Concentration[b] | Treatment | 1 | 143017 | 143017 | 15.13 | 0.000315 |
| | Concentration | 3 | 3827240 | 1275747 | 134.95 | < 2e-16 |
| | Strain | 2 | 1773619 | 886809 | 93.81 | < 2e-16 |
| | Treatment:Concentration | 3 | 1653257 | 551086 | 58.30 | 7.17e-16 |
| | Treatment:Strain | 2 | 1784833 | 892417 | 94.40 | < 2e-16 |
| | Concentration:Strain | 6 | 9157936 | 1526323 | 161.46 | < 2e-16 |
| | Treatment:Concentration:Strain | 6 | 5517365 | 919561 | 97.28 | < 2e-16 |
| | Residuals | 47 | 444303 | 9453 | | |
| Accumulation LOM Source (Carbon)[c] | Treatment | 1 | 5012379 | 5012379 | 56.270 | 8.71e-09 |
| | Carbon | 2 | 2018650 | 1009325 | 11.331 | 0.000161 |
| | Strain | 2 | 18059959 | 9029980 | 101.373 | 2.75e-15 |
| | Treatment:Carbon | 2 | 722775 | 361387 | 4.057 | 0.026023 |
| | Treatment:Strain | 2 | 21359893 | 10679946 | 119.896 | < 2e-16 |
| | Carbon:Strain | 4 | 1172539 | 293135 | 3.291 | 0.021667 |
| | Treatment:Carbon:Strain | 4 | 1423517 | 355879 | 3.995 | 0.008985 |
| | Residuals | 35 | 3117682 | 89077 | | |
| Rates Concentration[b] | Treatment | 1 | 134.5 | 134.54 | 425.61 | <2e-16 |
| | Concentration | 3 | 219.1 | 73.02 | 231.00 | <2e-16 |
| | Strain | 2 | 108.7 | 54.36 | 171.96 | <2e-16 |
| | Treatment:Concentration | 3 | 92.0 | 30.68 | 97.06 | <2e-16 |
| | Treatment:Strain | 2 | 125.0 | 62.48 | 197.66 | <2e-16 |
| | Concentration:Strain | 6 | 462.6 | 77.09 | 243.89 | <2e-16 |
| | Treatment:Concentration:Strain | 6 | 291.5 | 48.59 | 153.71 | <2e-16 |
| | Residuals | 47 | 14.9 | 0.32 | | |
| Rates LOM Source (Carbon)[c] | Treatment | 2 | 1257.5 | 628.7 | 907.40 | < 2e-16 |
| | Carbon | 2 | 88.4 | 44.2 | 63.77 | 1.02e-14 |
| | Strain | 2 | 645.5 | 322.8 | 465.82 | < 2e-16 |
| | Treatment:Carbon | 4 | 72.2 | 18.0 | 26.03 | 7.06e-12 |
| | Treatment:Strain | 4 | 1357.0 | 339.2 | 489.60 | < 2e-16 |
| | Carbon:Strain | 4 | 37.2 | 9.3 | 13.41 | 1.41e-07 |
| | Treatment:Carbon:Strain | 8 | 74.0 | 9.2 | 13.35 | 2.91e-10 |
| | Residuals | 52 | 36.0 | 0.7 | | |

[a]Due to the unbalanced nature of the respirometer experimental design two three-way ANOVAs were used to analyze the differences between mix and composite in terms of $CO_2$ accumulation and production rates. The two ANOVA models used tested whether the independent variables of inoculum, treatment, and either LOM source or concentration interacted to affect $CO_2$ accumulation or production rates.

[b]The independent variables assessed in this ANOVA model were inoculum (or strain), concentration, and treatment.

[c]The independent variables assessed in this ANOVA model were inoculum (or strain), LOM source (or carbon), and treatment.

**Table S6. Average Rate of $CO_2$ production (μg/h) for experiments shown in Figures 2 and 4.**

| Treatment | LOM | NOM[a] | MIX |
|---|---|---|---|
| SE45<br>400 μM-C Acetate | 6.29±1.85 | 0.98±0.21 | 5.65±1.68 |
| SE45<br>1 μM-C Casamino Acids | -1.09±0.80 | 0.98±0.21 | 1.46±0.03 |
| SE45<br>4 μM-C Casamino Acids | 0.84±0.08 | 1.99±0.41 | 1.61±0.51 |
| SE45<br>40 μM-C Casamino Acids | 2.90±0.44 | 1.99±0.41 | 695.02±358.06 |
| SE45<br>400 μM-C Casamino Acids | 6.38±0.07 | 1.99±0.41 | 3.44±0.71 |
| SE45<br>400 μM-C Coumarate | 6.06±0.29 | 0.98±0.21 | 7.68±2.06 |
| E-37<br>400 μM-C Acetate | 3.18±0.02 | 3.46±0.97 | 27.85±0.41 |
| E-37<br>1 μM-C Casamino Acids | -0.26±0.27 | 2.05±0.23 | 2.57±0.77 |
| E-37<br>4 μM-C Casamino Acids | -0.65±0.55 | 2.05±0.23 | 1.27±0.34 |
| E-37<br>40 μM-C Casamino Acids | -0.44±0.44 | 2.05±0.23 | 2.34±0.08 |
| E-37<br>400 μM-C Casamino Acids | 1.74±1.07 | 3.46±0.97 | 20.66±0.15 |
| E-37<br>400 μM-C Coumarate | 8.40±0.73 | 3.46±0.97 | 23.64±0.81 |
| Constructed Community<br>400 μM-C Acetate | 4.69±0.27 | 3.35±0.29 | 5.80 ±0.11 |
| Constructed Community<br>1 μM-C Casamino Acids | 2.45±0.15 | 3.19±0.07 | 4.29±0.22 |
| Constructed Community<br>4 μM-C Casamino Acids | 6.29±0.29 | 3.19±0.07 | 9.14±0.12 |
| Constructed Community<br>40 μM-C Casamino Acids | 8.90±0.07 | 3.19±0.07 | 6.12±0.38 |
| Constructed Community<br>400 μM-C Casamino Acids | 1.66±0.17 | 3.35±0.29 | 3.21±0.35 |
| Constructed Community<br>400 μM-C Coumarate | 9.00±0.94 | 3.35±0.29 | 9.10±0.22 |

[a] Samples were processed in batch. Variation in NOM alone values represents run-to-run variation as these controls were included in every batch run.

**Table S7. Probability values[a] for differences between respiration rates (μg/h) of LOM and mix.**

| LOM Source | LOM Concentration[b] 1 μM-C | 4 μM-C | 40 μM-C | 400 μM-C |
|---|---|---|---|---|
| **Acetate** | Not Measured | Not Measured | Not Measured | SE45: $p < 0.50$<br>E-37: $\boldsymbol{p < 0.001}$<br>Community: $p < 0.20$ |
| **Casamino Acids** | SE45: $\boldsymbol{p < 0.001}$<br>E-37: $\boldsymbol{p < 0.001}$<br>Community: $\boldsymbol{p < 0.001}$ | SE45: $p < 0.15$<br>E-37: $\boldsymbol{p < 0.001}$<br>Community: $\boldsymbol{p < 0.001}$ | SE45: $\boldsymbol{p < 0.001}$<br>E-37: $\boldsymbol{p < 0.001}$<br>Community: $p < 0.001$ | SE45: $p < 0.001$<br>E-37: $\boldsymbol{p < 0.001}$<br>Community: $p < 0.10$ |
| **Coumarate** | Not Measured | Not Measured | Not Measured | SE45: $\boldsymbol{p < 0.05}$<br>E-37: $\boldsymbol{p < 0.001}$<br>Community: $p < 0.95$ |

[a] Due to the unbalanced nature of the respirometer experimental design two three-way ANOVAs were used to analyze the differences between mix and LOM in terms of $CO_2$ production rates. The two ANOVA models used tested whether the independent variables of inoculum, treatment, and either LOM source or concentration interacted to affect $CO_2$ accumulation. As the 400 µM-C Casamino Acids accumulation was analyzed in both ANOVAs, the higher of the two resulting p-value from the post hoc test were used to determine significance. *p*-values are adjusted to correct for the false discovery rate using the Benjamini-Hochberg correction.

**Supplemental Table S8. ANOVA Tables for each three-way ANOVA performed for the constructed community cell density by day.**

| Day | ANOVA Table | | | | | |
|---|---|---|---|---|---|---|
| 1 | | Df | Sum Sq | Mean Sq | F value | Pr(>F) |
| | Treatment | 1 | 1.185e+11 | 1.185e+11 | 0.017 | 0.89602 |
| | Concentration | 3 | 1.616e+16 | 5.386e+15 | 779.351 | < 2e-16 |
| | Carbon | 3 | 5.357e+15 | 1.786e+15 | 258.377 | < 2e-16 |
| | Treatment:Concentration | 3 | 8.469e+13 | 2.823e+13 | 4.085 | 0.00836 |
| | Treatment:Carbon | 3 | 4.262e+13 | 1.421e+13 | 2.056 | 0.10956 |
| | Concentration:Carbon | 9 | 1.266e+16 | 1.407e+15 | 203.509 | < 2e-16 |
| | Treatment:Concentration:Carbon | 9 | 9.517e+13 | 1.057e+13 | 1.530 | 0.14447 |
| | Residuals | 124 | 8.570e+14 | 6.911e+12 | | |
| 2 | | Df | Sum Sq | Mean Sq | F value | Pr(>F) |
| | Treatment | 1 | 4.666e+13 | 4.666e+13 | 2.004 | 0.160 |
| | Concentration | 3 | 1.809e+16 | 6.032e+15 | 259.014 | <2e-16 |
| | Carbon | 3 | 3.011e+15 | 1.004e+15 | 43.106 | <2e-16 |
| | Treatment:Concentration | 3 | 1.146e+14 | 3.819e+13 | 1.640 | 0.185 |
| | Treatment:Carbon | 3 | 4.151e+13 | 1.384e+13 | 0.594 | 0.620 |
| | Concentration:Carbon | 9 | 8.992e+15 | 9.991e+14 | 42.903 | <2e-16 |
| | Treatment:Concentration:Carbon | 9 | 1.691e+14 | 1.879e+13 | 0.807 | 0.611 |
| | Residuals | 101 | 2.352e+15 | 2.329e+13 | | |
| 4 | | Df | Sum Sq | Mean Sq | F value | Pr(>F) |
| | Treatment | 1 | 4.026e+10 | 4.026e+10 | 0.003 | 0.95598 |
| | Concentration | 3 | 8.482e+15 | 2.827e+15 | 214.882 | < 2e-16 |
| | Carbon | 3 | 3.295e+15 | 1.098e+15 | 83.482 | < 2e-16 |
| | Treatment:Concentration | 3 | 6.241e+13 | 2.080e+13 | 1.581 | 0.19732 |
| | Treatment:Carbon | 3 | 1.543e+14 | 5.142e+13 | 3.908 | 0.01045 |
| | Concentration:Carbon | 9 | 8.542e+15 | 9.491e+14 | 72.132 | < 2e-16 |
| | Treatment:Concentration:Carbon | 9 | 3.956e+14 | 4.395e+13 | 3.340 | 0.00108 |
| | Residuals | 125 | 1.645e+15 | 1.316e+13 | | |
| 7 | | Df | Sum Sq | Mean Sq | F value | Pr(>F) |
| | Treatment | 1 | 5.033e+13 | 5.033e+13 | 5.902 | 0.016597 |
| | Concentration | 3 | 4.782e+15 | 1.594e+15 | 186.894 | < 2e-16 |
| | Carbon | 3 | 1.219e+15 | 4.063e+14 | 47.648 | < 2e-16 |
| | Treatment:Concentration | 3 | 1.778e+14 | 5.925e+13 | 6.948 | 0.000236 |
| | Treatment:Carbon | 3 | 1.985e+13 | 6.617e+12 | 0.776 | 0.509620 |
| | Concentration:Carbon | 9 | 3.138e+15 | 3.487e+14 | 40.884 | < 2e-16 |
| | Treatment:Concentration:Carbon | 9 | 8.561e+13 | 9.513e+12 | 1.115 | 0.357067 |
| | Residuals | 121 | 1.032e+15 | 8.528e+12 | | |
| 10 | | Df | Sum Sq | Mean Sq | F value | Pr(>F) |
| | Treatment | 1 | 1.178e+14 | 1.178e+14 | 12.658 | 0.000532 |
| | Concentration | 3 | 6.171e+15 | 2.057e+15 | 221.110 | < 2e-16 |
| | Carbon | 3 | 2.062e+15 | 6.874e+14 | 73.895 | < 2e-16 |
| | Treatment:Concentration | 3 | 5.647e+14 | 1.882e+14 | 20.234 | 1.00e-10 |
| | Treatment:Carbon | 3 | 2.959e+14 | 9.865e+13 | 10.604 | 2.99e-06 |
| | Concentration:Carbon | 9 | 5.634e+15 | 6.260e+14 | 67.292 | < 2e-16 |
| | Treatment:Concentration:Carbon | 9 | 6.062e+14 | 6.736e+13 | 7.241 | 2.14e-08 |
| | Residuals | 123 | 1.144e+15 | 9.303e+12 | | |
| 14 | | Df | Sum Sq | Mean Sq | F value | Pr(>F) |
| | Treatment | 1 | 7.619e+13 | 7.619e+13 | 15.55 | 0.000144 |
| | Concentration | 3 | 4.007e+15 | 1.336e+15 | 272.65 | < 2e-16 |
| | Carbon | 3 | 1.438e+15 | 4.795e+14 | 97.88 | < 2e-16 |
| | Treatment:Concentration | 3 | 3.158e+14 | 1.053e+14 | 21.48 | 6.01e-11 |
| | Treatment:Carbon | 3 | 1.636e+14 | 5.453e+13 | 11.13 | 2.07e-06 |
| | Concentration:Carbon | 9 | 3.337e+15 | 3.708e+14 | 75.69 | < 2e-16 |
| | Treatment:Concentration:Carbon | 9 | 4.453e+14 | 4.948e+13 | 10.10 | 4.33e-11 |
| | Residuals | 106 | 5.193e+14 | 4.899e+12 | | |

**Supplemental Table S9. ANOVA tables for each three-way ANOVA performed for constructed community alpha diversity estimates by day.**

| Day | ANOVA Table | | | | | |
|---|---|---|---|---|---|---|
| 1 | | Df | Sum Sq | Mean Sq | F value | Pr(>F) |
| | Treatment | 1 | 1.89 | 1.89 | 8.115 | 0.005141 |
| | Concentration | 3 | 110.94 | 36.98 | 158.782 | < 2e-16 |
| | Carbon | 3 | 28.73 | 9.58 | 41.123 | < 2e-16 |
| | Treatment:Concentration | 3 | 0.64 | 0.21 | 0.922 | 0.432564 |
| | Treatment:Carbon | 3 | 1.73 | 0.58 | 2.479 | 0.064284 |
| | Concentration:Carbon | 9 | 16.19 | 1.80 | 7.725 | 5.8e-09 |
| | Treatment:Concentration:Carbon | 9 | 8.51 | 0.95 | 4.062 | 0.000138 |
| | Residuals | 124 | 28.88 | 0.23 | | |
| 2 | | Df | Sum Sq | Mean Sq | F value | Pr(>F) |
| | Treatment | 1 | 0.39 | 0.39 | 1.037 | 0.310547 |
| | Concentration | 3 | 125.41 | 41.80 | 110.947 | < 2e-16 |
| | Carbon | 3 | 13.83 | 4.61 | 12.235 | 4.71e-07 |
| | Treatment:Concentration | 3 | 1.60 | 0.53 | 1.418 | 0.240679 |
| | Treatment:Carbon | 3 | 0.58 | 0.19 | 0.516 | 0.671937 |
| | Concentration:Carbon | 9 | 13.47 | 1.50 | 3.973 | 0.000182 |
| | Treatment:Concentration:Carbon | 9 | 3.29 | 0.37 | 0.971 | 0.467301 |
| | Residuals | 122 | 45.97 | 0.38 | | |
| 4 | | Df | Sum Sq | Mean Sq | F value | Pr(>F) |
| | Treatment | 1 | 0.00 | 0.003 | 0.015 | 0.904 |
| | Concentration | 3 | 59.15 | 19.718 | 103.707 | < 2e-16 |
| | Carbon | 3 | 7.13 | 2.377 | 12.500 | 3.39e-07 |
| | Treatment:Concentration | 3 | 4.75 | 1.583 | 8.328 | 4.34e-05 |
| | Treatment:Carbon | 3 | 1.05 | 0.348 | 1.832 | 0.145 |
| | Concentration:Carbon | 9 | 10.01 | 1.112 | 5.851 | 8.91e-07 |
| | Treatment:Concentration:Carbon | 9 | 9.34 | 1.037 | 5.456 | 2.67e-06 |
| | Residuals | 124 | 23.58 | 0.190 | | |
| 7 | | Df | Sum Sq | Mean Sq | F value | Pr(>F) |
| | Treatment | 1 | 0.37 | 0.365 | 3.344 | 0.0699 |
| | Concentration | 3 | 39.20 | 13.066 | 119.573 | < 2e-16 |
| | Carbon | 3 | 5.25 | 1.752 | 16.030 | 7.74e-09 |
| | Treatment:Concentration | 3 | 1.19 | 0.398 | 3.639 | 0.0148 |
| | Treatment:Carbon | 3 | 1.04 | 0.348 | 3.186 | 0.0263 |
| | Concentration:Carbon | 9 | 6.13 | 0.681 | 6.232 | 3.36e-07 |
| | Treatment:Concentration:Carbon | 9 | 4.19 | 0.465 | 4.258 | 8.15e-05 |
| | Residuals | 121 | 13.22 | 0.109 | | |
| 10 | | Df | Sum Sq | Mean Sq | F value | Pr(>F) |
| | Treatment | 1 | 10.731 | 10.731 | 125.149 | < 2e-16 |
| | Concentration | 3 | 30.257 | 10.086 | 117.623 | < 2e-16 |
| | Carbon | 3 | 7.102 | 2.367 | 27.608 | 1.01e-13 |
| | Treatment:Concentration | 3 | 1.474 | 0.491 | 5.730 | 0.00105 |
| | Treatment:Carbon | 3 | 0.191 | 0.064 | 0.743 | 0.52814 |
| | Concentration:Carbon | 9 | 4.403 | 0.489 | 5.705 | 1.36e-06 |
| | Treatment:Concentration:Carbon | 9 | 3.389 | 0.377 | 4.391 | 5.46e-05 |
| | Residuals | 123 | 10.547 | 0.086 | | |
| 14 | | Df | Sum Sq | Mean Sq | F value | Pr(>F) |
| | Treatment | 1 | 19.789 | 19.789 | 215.291 | < 2e-16 |
| | Concentration | 3 | 17.150 | 5.717 | 62.191 | < 2e-16 |
| | Carbon | 3 | 25.118 | 8.373 | 91.088 | < 2e-16 |
| | Treatment:Concentration | 3 | 5.414 | 1.805 | 19.635 | 1.88e-10 |
| | Treatment:Carbon | 3 | 1.483 | 0.494 | 5.379 | 0.001640 |
| | Concentration:Carbon | 9 | 10.376 | 1.153 | 12.543 | 6.26e-14 |
| | Treatment:Concentration:Carbon | 9 | 2.833 | 0.315 | 3.425 | 0.000867 |
| | Residuals | 122 | 11.214 | 0.092 | | |

### Supplemental Figures

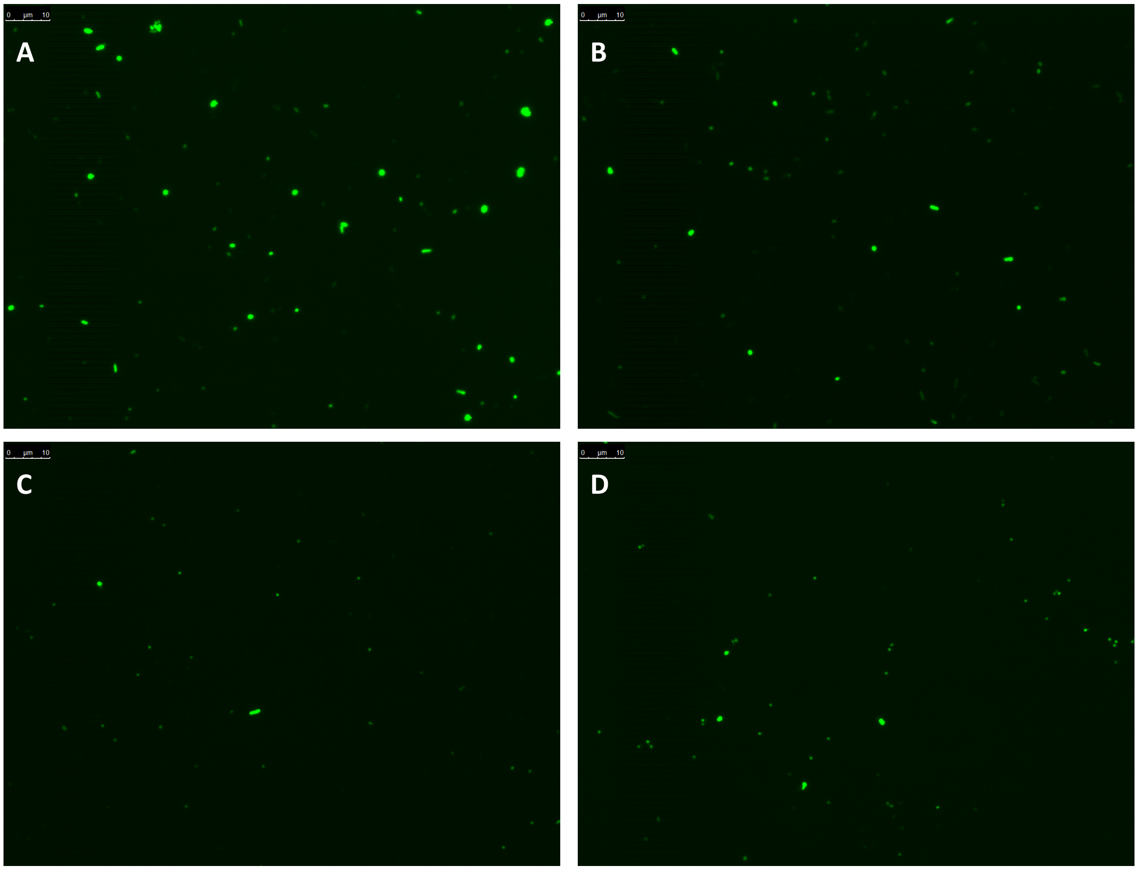


**Supplemental Figure 1.** Microscopic images of representative constructed community cultures grown on (A & B) 400 uM casamino acids + NOM on Day 3; replicate cultures and (C & D) on only 400 uM casamino acids on Day 6. Cultures were stained with DAPI and collected on a 0.22- µm polycarbonate filter. A scale bar is provided in the upper left-hand corner of each image.

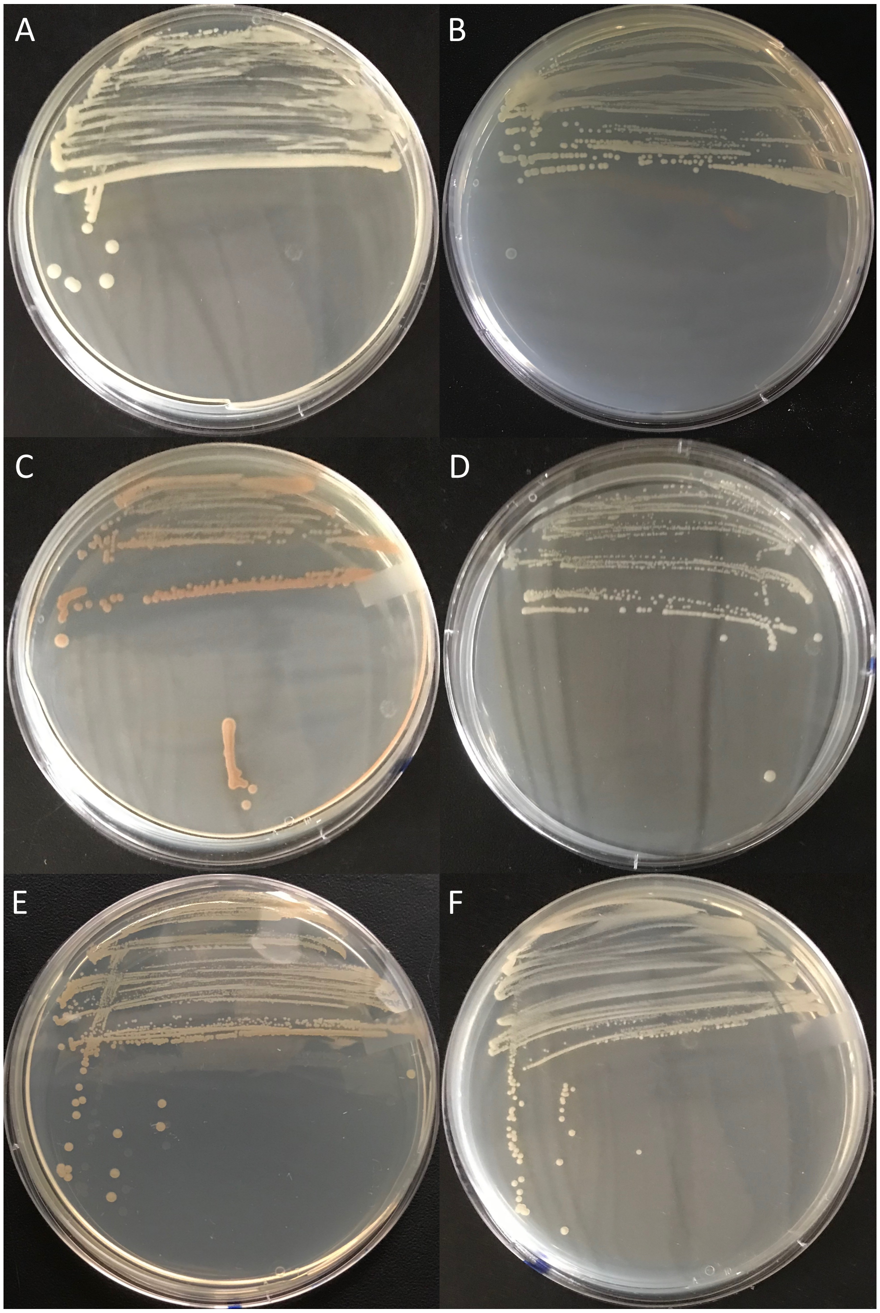


**Supplemental Figure 2. Colony morphologies of the bacterial isolates used in the constructed community.** A. *Citreicella* sp. SE45 B. *Phaeobacter* sp. Y4I C. *Roseovarius nubinhibens* ISM D. *Sagittula stellata* E-37 E. *Sulfitobacter* sp. EE-36 F. *Sulfitobacter* sp. NAS-14.1

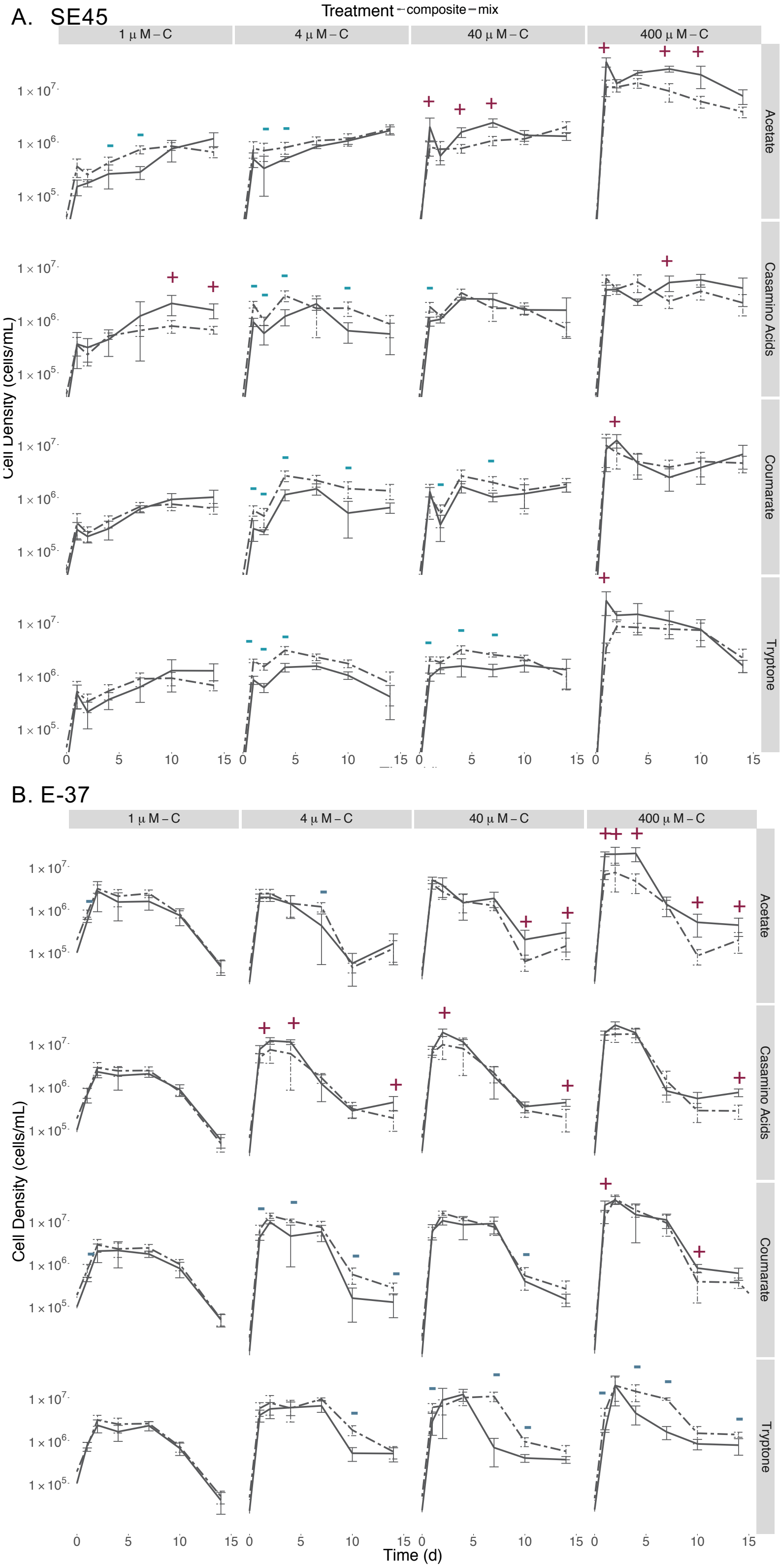

A. SE45
Treatment composite mix
1 µ M – C
4 µ M – C
40 µ M – C
400 µ M – C
Acetate
Casamino Acids
Coumarate
Tryptone
Cell Density (cells/mL)
1 × 10^7
1 × 10^6
1 × 10^5
0
5
10
15
B. E-37
Time (d)

**Supplemental Figure S3. Viable counts for monocultures of (A) SE45 and (B) E-37 in composite (dashed line) and mix (black line) treatments.**

Points represent the mean (n=3-5); error bars represent one standard deviation from the mean. Red plus signs indicate a significant synergistic interactive effect ($p < 0.05$), blue minus signs indicate a significant antagonistic interactive effect ($p < 0.05$). Seeding densities for SE45 and E-37 and six-member constructed community were $1.51 \times 10^4$ CFU/mL ($\pm 5.1 \times 10^3$) and $4.23 \times 10^4$ CFU/mL ($\pm 9 \times 10^3$). The composite treatment is the sum of results of the LOM and NOM treatments.

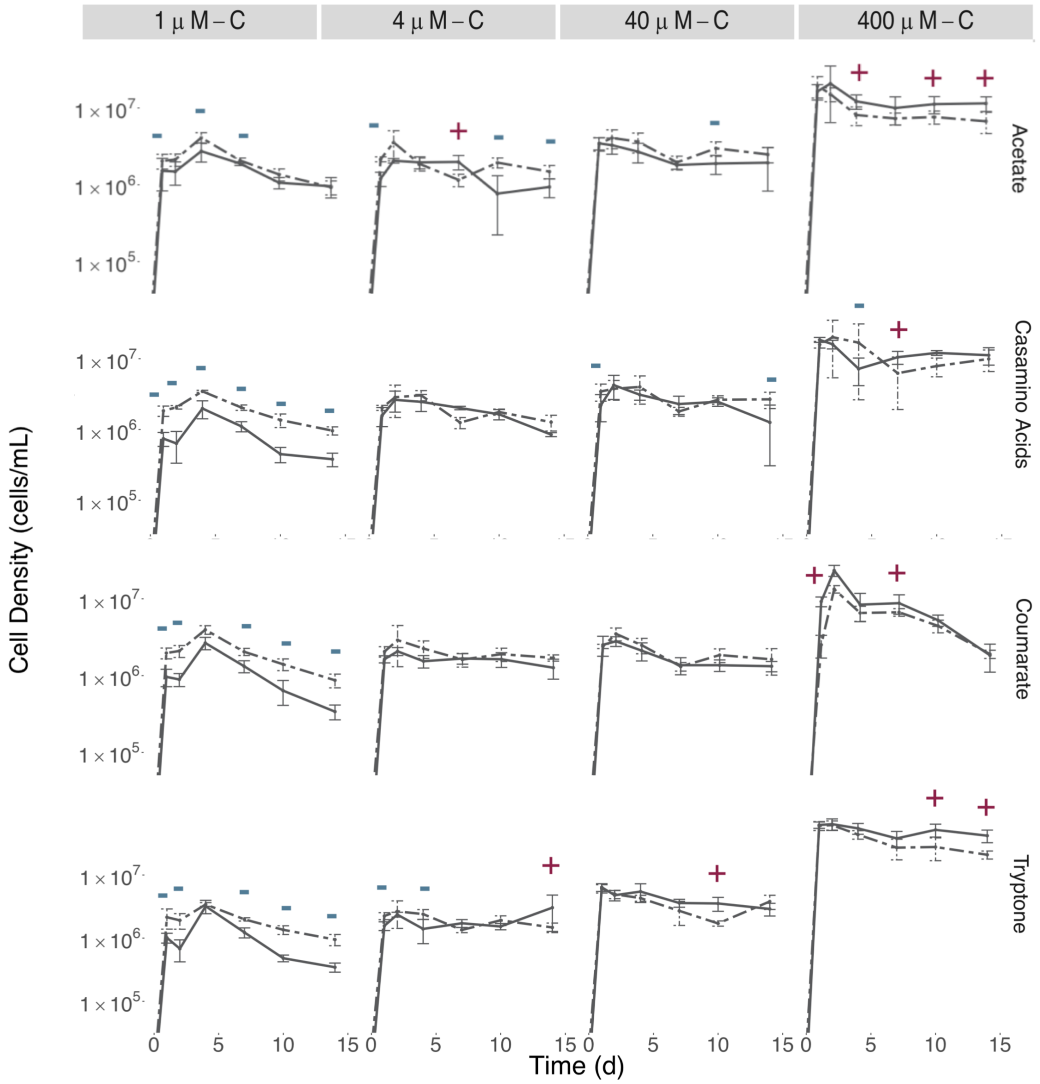


**Supplemental Figure S4. Viable counts for constructed communities in composite (dashed and mix (black line) treatments.**

Points represent the mean (n=3-5); error bars represent one standard deviation from the mean. Red plus signs indicate a significant synergistic interactive effect ($p < 0.05$), blue minus signs indicate a significant antagonistic interactive effect ($p < 0.05$). Seeding densities for six-member constructed community was 7.01 x $10^3$ CFU/mL (±2.6 x $10^3$), respectively. The composite treatment is the sum of results of the LOM and NOM treatments.

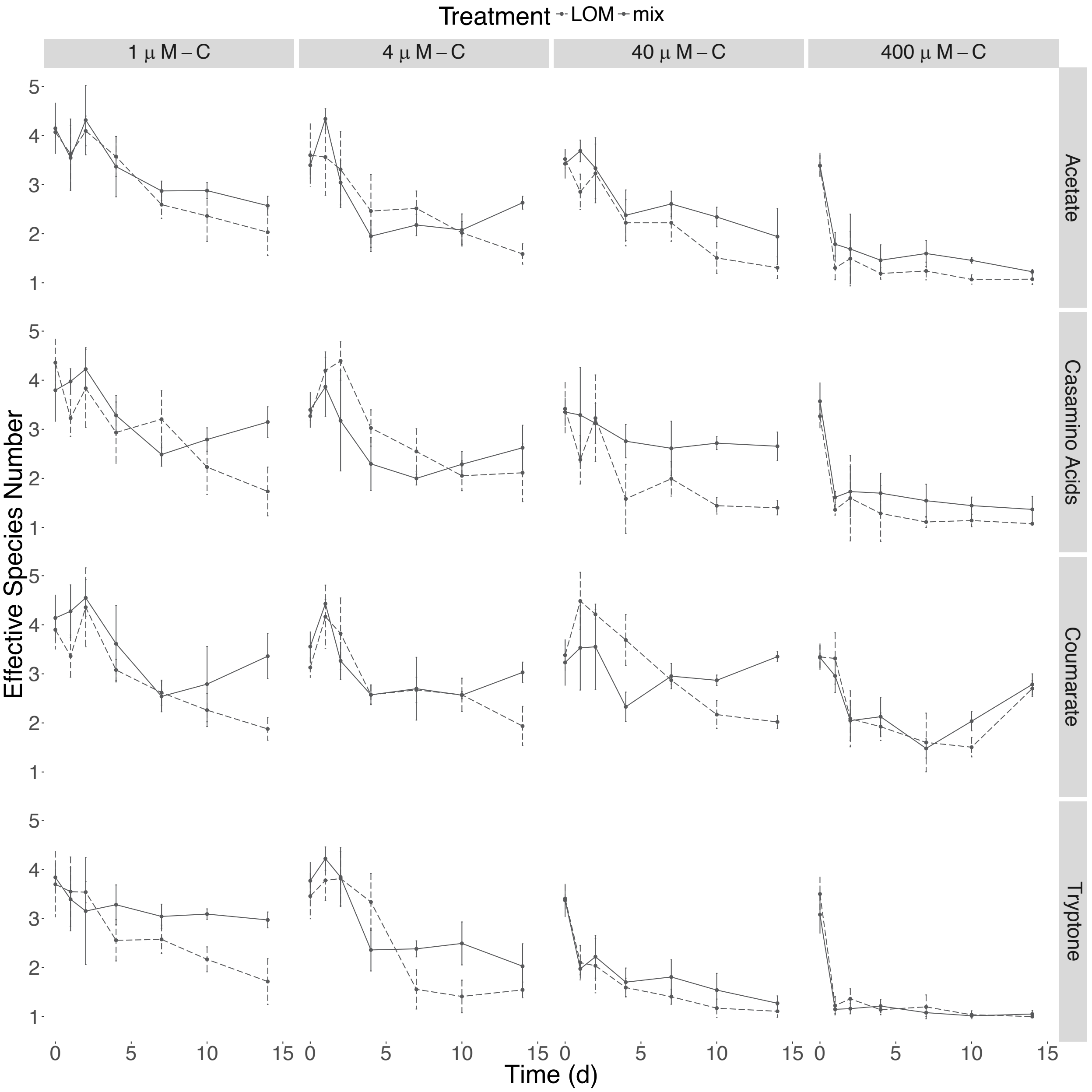


**Supplemental Figure S5. Alpha diversity of constructed community under mixed substrate conditions.**

Shannon entropy was calculated for each replicate at each time point. Shannon entropy was converted to Hill numbers or effective species number. Points represent the average of 3-5 replicates, and error bars represent one standard deviation from the mean. Dashed lines represent the LOM cultures while the solid lines represent the mixed carbon cultures.

**A.**

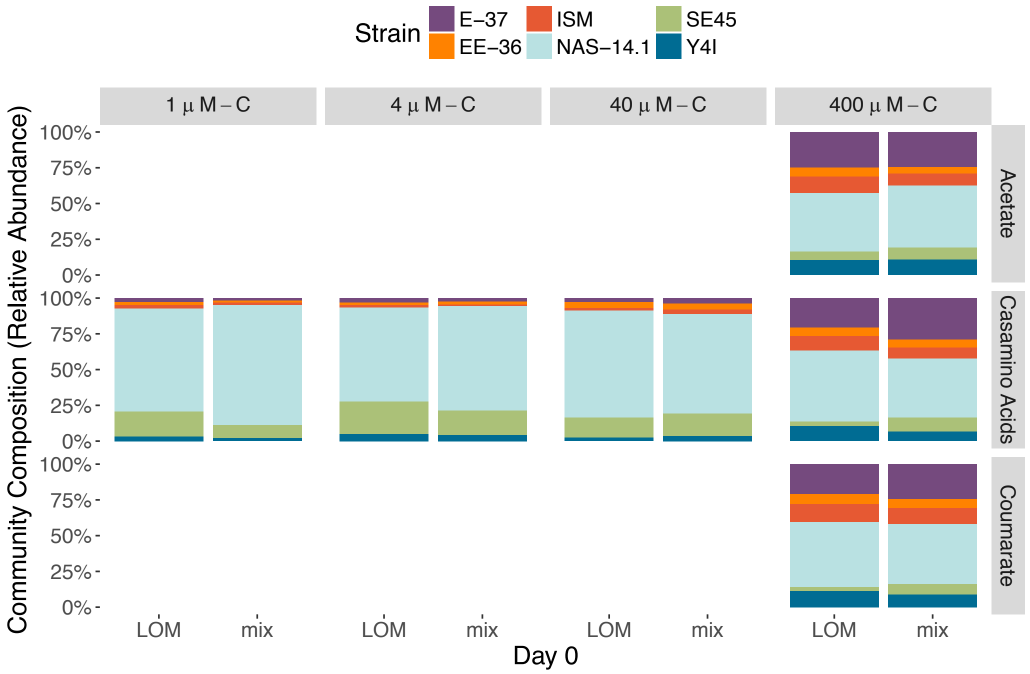

Strain
E–37
EE–36
ISM
NAS–14.1
SE45
Y4I
1 µ M – C
4 µ M – C
40 µ M – C
400 µ M – C
Acetate
Casamino Acids
Coumarate
Community Composition (Relative Abundance)
100%
75%
50%
25%
0%
LOM
mix
Day 0


**B.**

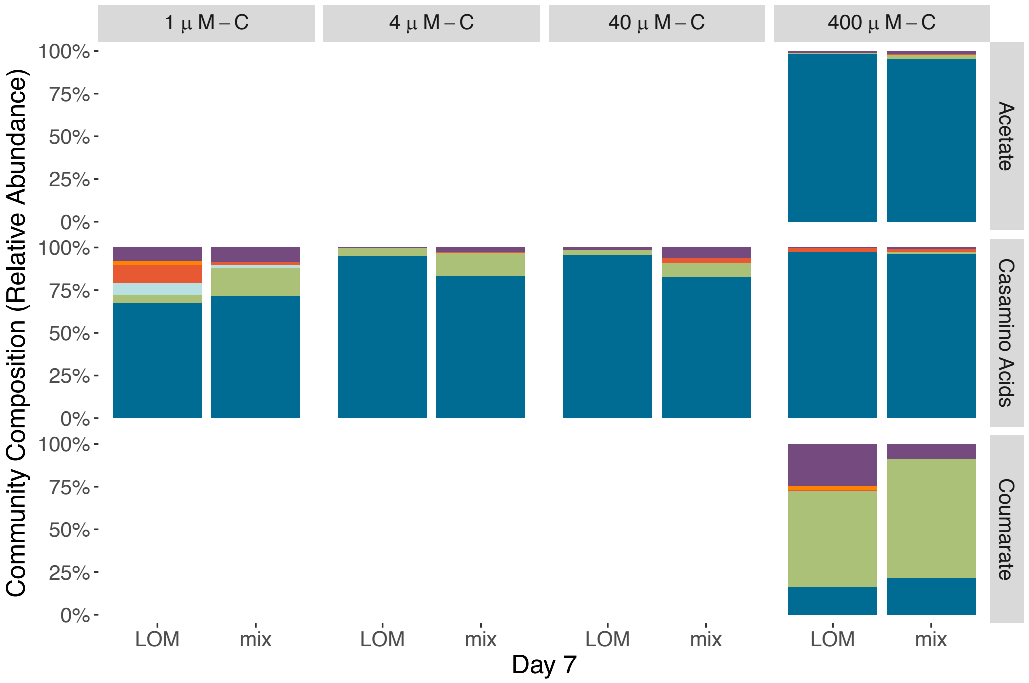

1 µ M – C
4 µ M – C
40 µ M – C
400 µ M – C
Acetate
Casamino Acids
Coumarate
Community Composition (Relative Abundance)
100%
75%
50%
25%
0%
LOM
mix
Day 7

C.

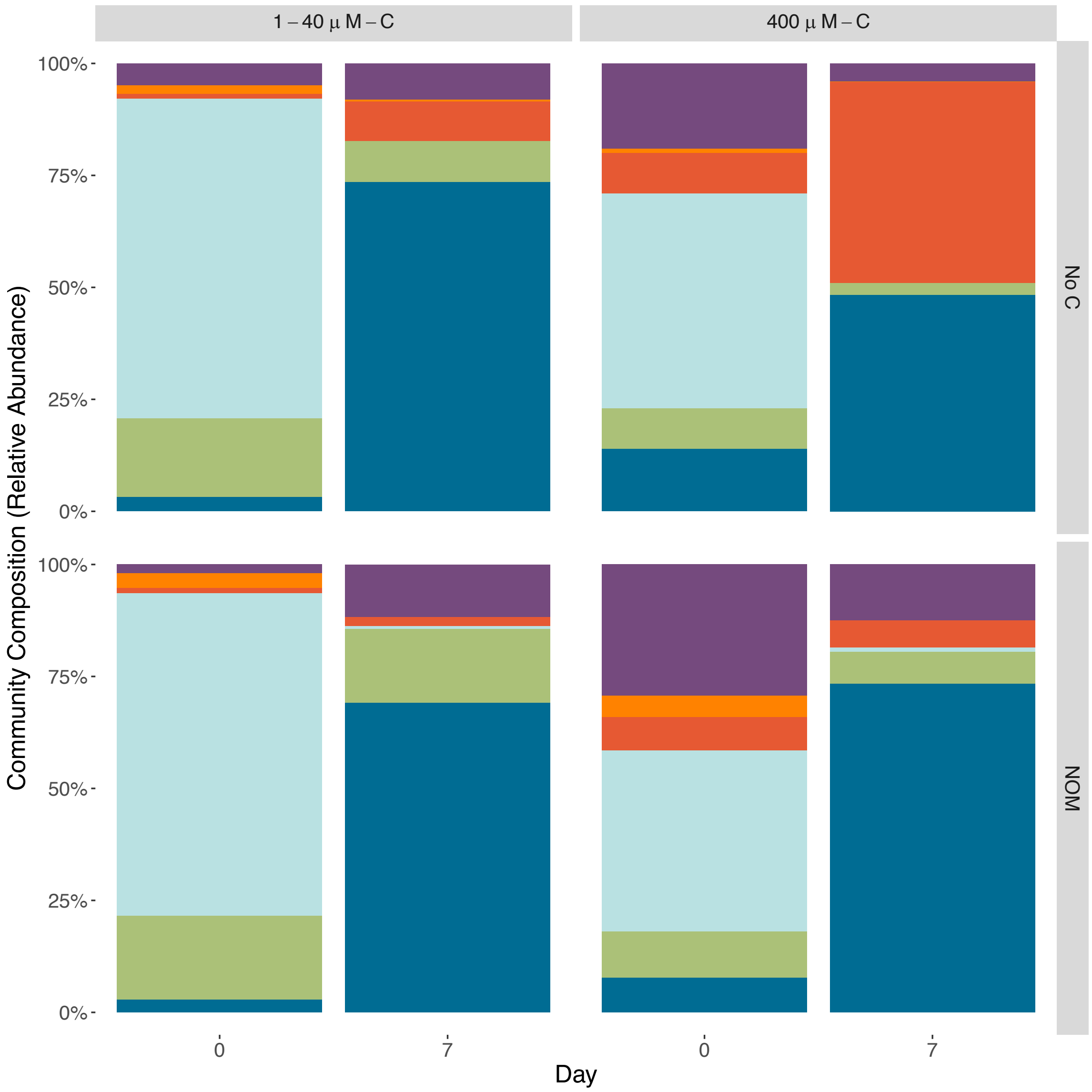


**Supplemental Figure S6. The community composition of the respirometer incubations for the LOM and mix treatments at Day 0 (A) and Day 7. (B)** The community composition for the NOM and No C treatments are present in **panel C**. The NOM and No C treatments for the first incubation which included the low concentrations of Casamino Acids (1, 4, and 40 uM) are in the left panel while the community composition for the NOM and No C treatments for the high concentrations of labile carbon are in the right panel.

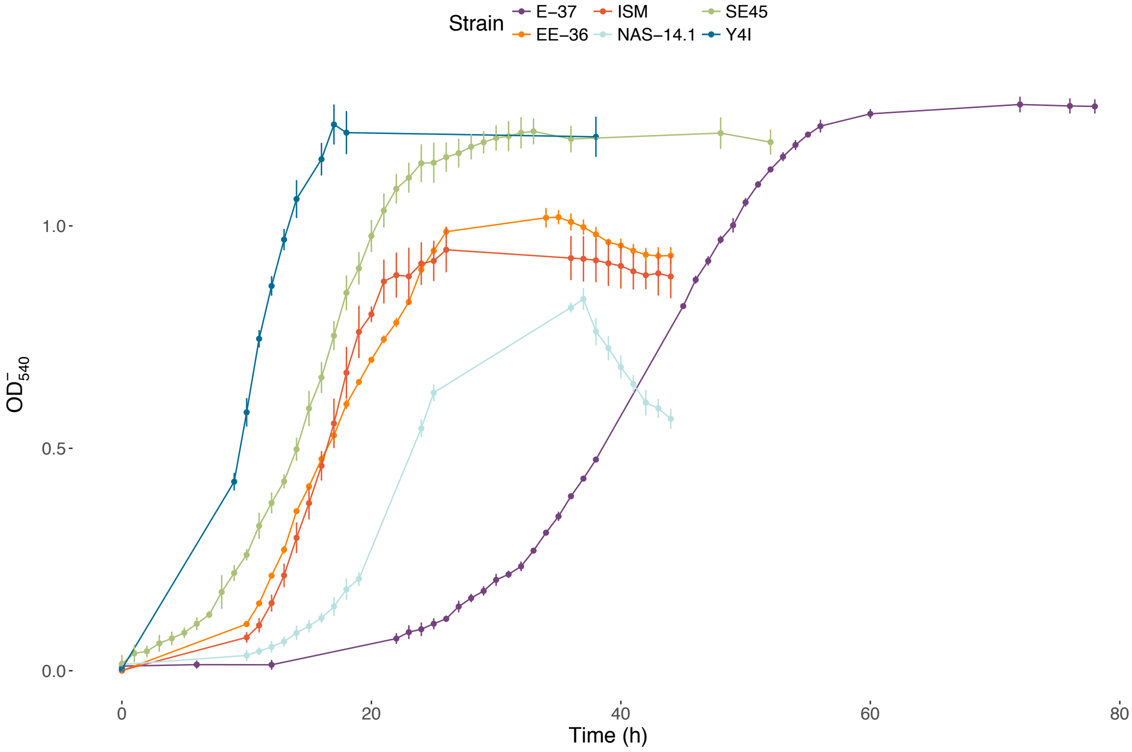


**Supplemental Figure S7 Growth curves of the strains used in this study when grown as monocultures in minimal media supplemented with 2 mM-C tryptone.** Points represent the mean of three biological replicates and error bars represent one standard deviation from the mean.
